# Obsessive-Compulsive Tendencies Shift the Balance Between Competitive Neurocognitive Functions

**DOI:** 10.1101/2025.08.13.669948

**Authors:** Bianka Brezóczki, Teodóra Vékony, Bence Csaba Farkas, Flóra Hann, Dezső Németh

## Abstract

Theoretical models of Obsessive–Compulsive Disorder (OCD) emphasize that symptoms may arise from an imbalance between habitual and goal-directed processes, characterized by increased reliance on habitual behavior and reduced efficiency of goal-directed control. However, it remains unclear whether similar alterations appear at a more general functional level, beyond reward-driven mechanisms. The present study, therefore, investigated the relationship between statistical learning (SL), an implicit, reward-independent mechanism that supports the detection of environmental regularities and the formation of habitual behaviour, and cognitive flexibility, defined as the capacity to adapt behaviour and cognitive strategies to changing environmental demands. By adopting a dimensional approach to obsessive-compulsive (OC) tendencies in a university student sample, we aimed to clarify how continuous symptom variability relates to the interaction of these neurocognitive functions. A total of 404 participants completed an online experiment, including a probabilistic sequence learning task assessing SL and a card-sorting task measuring cognitive flexibility. Results revealed an antagonistic relationship between SL and cognitive flexibility. Importantly, this inverse association weakened as OC tendencies increased, suggesting that OC tendencies may alter the balance between automatic and goal-directed functions.

**Highlights:**

- CD theories propose an imbalance between automatic and goal-directed control systems.
- tested whether this imbalance emerges at a functional level, beyond reward-based learning.
- learning and cognitive flexibility showed an antagonistic relationship in a university student sample.
- inverse association weakened as obsessive–compulsive tendencies increased.
- C tendencies alter the interaction between automatic learning and executive control.

**Graphical Abstract:** 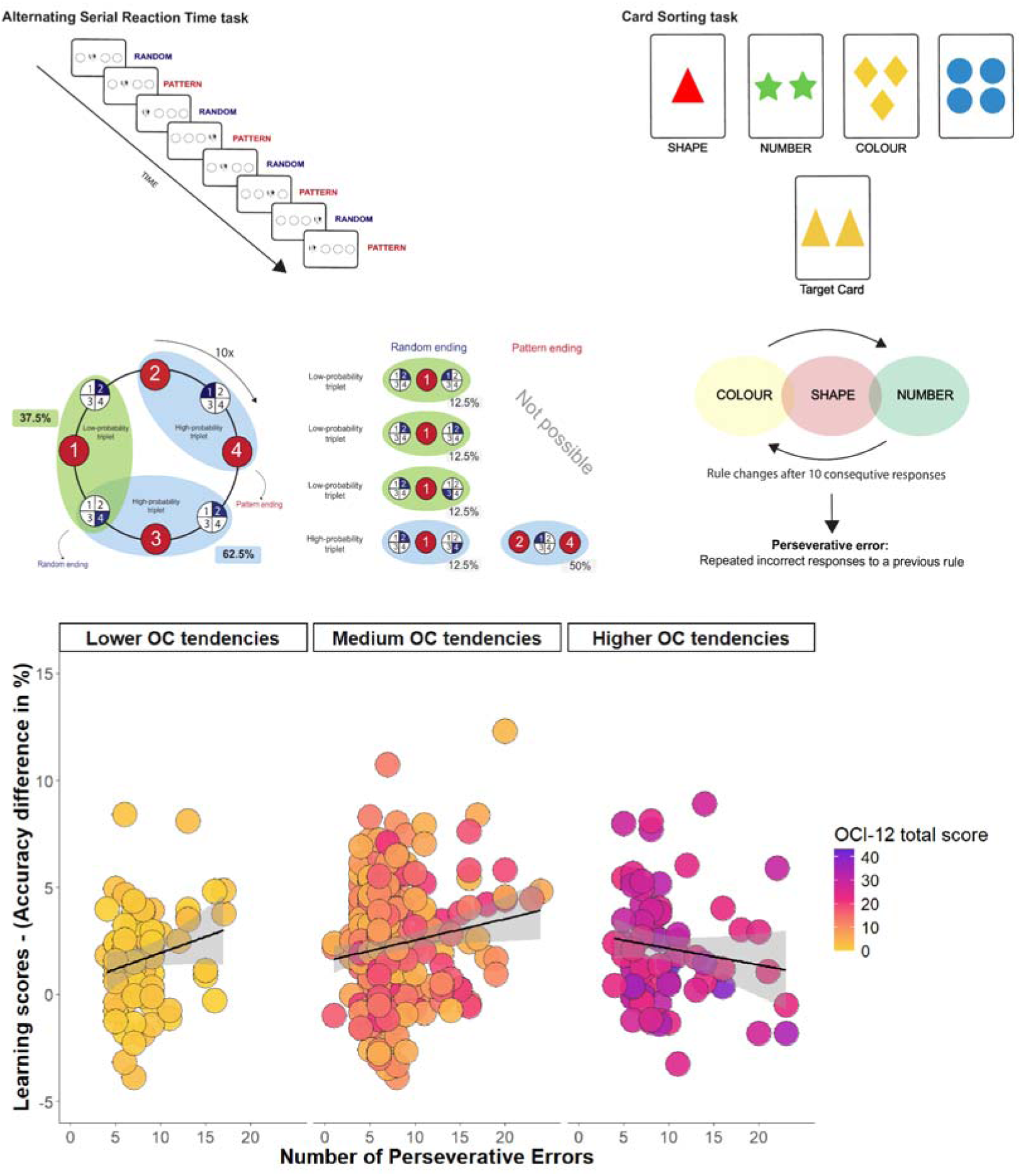

## Introduction

A core challenge in human cognition involves maintaining a dynamic interplay between efficient automatic and flexible goal-directed functions to support adaptive behavior. Goal-directed behaviour relies on a broad set of executive functions - including cognitive flexibility, inhibitory control, and working memory updating - that support the attainment of desired outcomes and avoiding undesired ones ^1–3^. In contrast, automatic behaviour depends on previously acquired associations and operates with reduced conscious engagement ^4–6^. Disruption of this interplay can lead to behavioral difficulties, as seen in conditions such as Obsessive-Compulsive Disorder (OCD), where automaticity tends to dominate over goal-directed control, resulting in rigid, repetitive mental (obsessions) and behavioral rituals (compulsions)^7–14^. Indeed, varying degrees of obsessions and compulsive acts, collectively referred to as obsessive-compulsive (OC) tendencies, are distributed along a continuum in the general population and, when persistent and sufficiently severe, culminate in a clinical presentation of OCD ^15–23^. Considering OC tendencies from a dimensional continuum provides a more precise and comprehensive identification of the neurocognitive mechanisms underlying OCD, effectively overcoming the limitations of traditional categorical diagnoses that often obscure the continuous and nuanced variability of these symptoms ^11,15,24–26^. Therefore, the present study aimed to delineate the role of the balance between automatic and goal-directed functions underlying maladaptive behaviours of OCD by assessing OC tendencies as a dimensional spectrum within the non-clinical university student population. Prominent theoretical models propose that the repetitive symptoms of OCD and transdiagnostic tendencies arise from diminished goal-directed control coupled with a corresponding dominance of habitual responding ^8,9,12,14,27–29^. Evidence for this framework has been driven largely by dual-task paradigms, which show that increasing cognitive demands can bias the expression of these processes within a given task ^8,9,12,14,27–29^. While these findings establish that controlled and automatized systems interact in a demand-sensitive manner, they are inherently constrained to within-task dynamics, leaving unresolved whether comparable patterns extend across tasks that differentially engage these processes ^8,9,30,32^. These approaches do not clearly distinguish between competing mechanistic accounts of this shift, whether it reflects increased reliance on habitual processes, reduced goal-directed control, or a disruption in the coordination between the two. Moreover, it remains unclear whether these imbalances are consistent across tasks or whether they primarily reflect context-dependent performance constraints within specific paradigms rather than more stable individual differences. The present study addresses this question by testing whether inter-individual variability in performance is systematically aligned across paradigms that place distinct demands on controlled and automatized processing, and whether such alignment relates to OC tendencies. By shifting the focus from within-task modulation to cross-task structure, this approach offers a complementary perspective on how variability in cognitive performance is organized across individuals, thereby refining current accounts of the mechanisms implicated in compulsivity. From a habitual perspective, a key limitation of current frameworks becomes apparent. Evidence from outcome devaluation and related supervision-based paradigms consistently points to enhanced habitual responding in OCD and compulsivity ^10,12,28^. Yet in these tasks, the balance between goal-directed and habitual control is tightly coupled to continuous external reward or feedback, making it difficult to isolate learning mechanisms from reward-driven performance. Consequently, understanding OC requires moving beyond feedback-dependent paradigms to probe the underlying, feedback-independent mechanisms ^21^. Addressing this question requires considering recent advances in learning theory, which emphasize the crucial role of unsupervised processes in driving neural plasticity and establishing the foundation for supervised learning ^30^. From this perspective, supervised learning emerges as a later process built upon earlier, reward-free, unsupervised mechanisms ^30^. Accordingly, to better understand the development and persistence of automatic, compulsive behavioral characteristics of OCD and OC tendencies, it is essential to disentangle feedback sensitivity and pure learning mechanisms ^21^ and to revisit the primary role of unsupervised learning and its interaction with prefrontal cortex-related functions.

One such unsupervised learning mechanism is statistical learning (SL), a fundamental function that supports the acquisition of skills and habits through the incidental extraction of novel probabilistic patterns from the environmental stream ^6,31–34^. SL in clinical OCD populations has shown both reduced and intact results (for review see: Brezóczki et al. ^35^), while individuals scoring on the dimensional OC scale demonstrate intact or even faster learning rates ^21^. This highlights the need to better understand how other neurocognitive functions contribute to such performance. Indeed, within the framework of interactive memory systems, neurocognitive functions such as SL rarely operate in isolation; instead, they engage in dynamic and often competitive interactions with goal-directed functions ^6,33,36–40^. Typically, increased reliance on one system corresponds with reduced engagement of the other, as demonstrated by the inverse relationship between SL and goal-directed functions. In this context, cognitive flexibility, defined as the capacity to shift behavior, strategies, or mental frameworks in response to changing environmental demands, is a particularly critical goal-directed function ^1–3^, as maladaptive, rigid, and repetitive behavioral patterns of OC may reflect its impairments ^41,42^. If OCD and OC tendencies modulate a disruption in the balance between automatic and goal-directed functions, then investigating their interaction outside the influence of supervision becomes particularly informative.

To provide a complementary perspective on the neurocognitive profile underlying OCD and OC tendencies, this study focused on SL and goal-directed function, specifically cognitive flexibility. Therefore, in the present study, we recruited 404 adult participants from university courses via an online experiment. SL was assessed using an unsupervised learning paradigm, the Alternating Serial Reaction Time (ASRT) task ^43,44^, which is a probabilistic learning task that lacks the supervision and reinforcement-based factors on SL performance^45^. Cognitive flexibility was operationalized as the number of perseverative errors on the Card Sorting Task ^46^, a widely accepted and extensively used measure of cognitive flexibility ^47–53^. OC tendencies were assessed dimensionally using the OCI-12 questionnaire ^54^. The present study aimed to uncover the interplay between SL and cognitive flexibility and to investigate the individual differences in their interaction among university students scoring on the dimensional OC measure. Based on existing literature, we hypothesized an antagonistic relationship between SL and cognitive flexibility, such that stronger performance on the ASRT task would be associated with weaker performance on the Card Sorting Task. Crucially, we further hypothesized that OC tendencies would modulate this interaction, with the relationship between SL and cognitive flexibility varying across different levels of OC tendencies and potentially diverging from a straightforward competitive pattern at higher levels of OC tendencies.

## Results

### How do OC tendencies modulate the interplay between cognitive flexibility and statistical learning? - Reaction Time

We performed a linear mixed-effects model on blockwise median RT of the ASRT task, with fixed effects for Time (Block 1-15), Triplet Type (high- vs. low-probability), OCI-12 score, Number of Perseverative errors of the Card Sorting Task, and all higher-order interactions. Random effects included participant-specific intercepts and a random slope for Time to account for individual variability. The statistical parameters underlying the effects reported above are summarized in Table 1, whereas the full set of statistical estimates (b, SE, 95% CI, t, df, and p values) is provided in Supplementary Table S1.

**Table 1.** Results of the Linear Mixed-Effects Model predicting blockwise median Reaction Time.

| Effect | *df* | *F* | *p* |
| --- | --- | --- | --- |
| Time | 1, 399.98 | 125.06 | < **.001** |
| Triplet Type | 1, 11297.07 | 169.21 | < **.001** |
| OCI-12 score | 1, 400.01 | 3.15 | .077 |
| Perseverative error | 1, 400.02 | 7.31 | **.007** |
| Time × Triplet Type | 1, 11297.10 | 10.42 | **.001** |
| Time × OCI-12 score | 1, 399.88 | 0.15 | .699 |
| Triplet Type × OCI-12 score | 1, 11297.11 | 0.00 | .926 |
| Time × Perseverative error | 1, 400.07 | 3.13 | .078 |
| Triplet Type × Perseverative error | 1, 11297.07 | 0.44 | .508 |
| OCI-12 score × Perseverative error | 1, 400.01 | 0.26 | .600 |
| Time × Triplet Type × OCI-12 score | 1, 11297.18 | 0.09 | .763 |
| Time × Triplet Type × Perseverative error | 1, 11297.1 | 0.92 | .338 |
| Time × OCI-12 score × Perseverative error | 1, 400.00 | 3.61 | .058 |
| Triplet Type × OCI-12 score × Perseverative error | 1, 11297.09 | 1.29 | .257 |
| Time × Triplet Type × OCI-12 score × Perseverative error | 1, 11297.15 | 0.67 | .412 |

The model reported a statistically significant main effect of Triplet Type in the ASRT task, indicating faster RTs for high-probability trials compared to low-probability trials, which evidences statistical learning during the task. The main effect of Time indicated a decrease in RTs throughout the task, demonstrating changes in visuomotor performance over time. Importantly, the interaction between Triplet Type and Time suggested that statistical learning developed gradually, as the difference between high- and low-probability trials increased over time.

Cognitive flexibility was significantly associated with overall RT performance, as reflected by a main effect of Perseverative errors on the CST, whereby a higher number of perseverative errors, indicative of reduced cognitive flexibility, predicted slower responses. However, the absence of statistically significant interactions involving Perseverative errors suggests that cognitive flexibility did not modulate visuomotor performance (Time × Perseverative error), or statistical learning (Triplet Type × Perseverative error), nor the trajectory of statistical learning (Time × Triplet Type × Perseverative error).

In contrast, OCI-12 scores showed no statistically significant main effect and did not interact with any of the task-related variables, indicating that OC tendencies were not significantly associated with baseline RT (main effect of OCI-12 score), cognitive flexibility (OCI-12 score × Perseverative error), visuomotor performance (Time × OCI-12 score), statistical learning (Triplet Type × OCI-12 score), the trajectory of statistical learning (Time × Triplet Type × OCI-12 score). Moreover, OCI-12 scores did not modulate the relationship between cognitive flexibility (Perseverative error × Triplet Type × OCI-12 score, see Figure 1) and the dynamics of statistical learning in the present study (Perseverative error × Time × Triplet Type × OCI-12 score).

**Figure 1.**
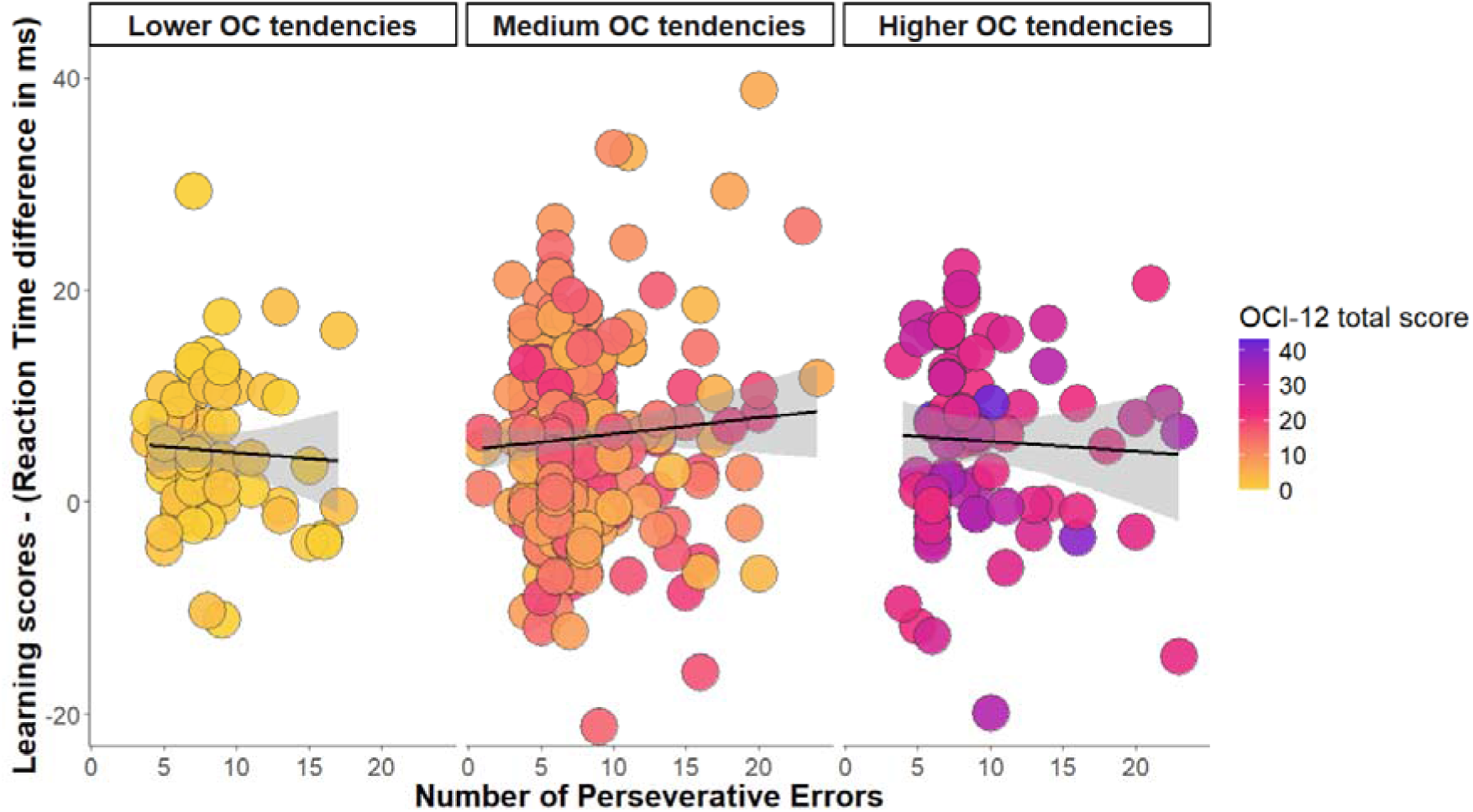
The figure illustrates the relationship between cognitive flexibility, as measured by the Number of Perseverative errors on the Card Sorting Task along the x-axis, and statistical learning performance along the y-axis, which is operationalized as the difference in reaction time between high- and low-probability triplets. Each point represents an individual participant, with color encoding their total OCI-12 score along a continuous gradient from low (yellow) through intermediate (pink) to high (purple) obsessive-compulsive tendencies. A linear regression line with 95% confidence intervals is overlaid in black. While the OCI-12 score was treated as a continuous variable in all statistical analyses, for visualization purposes, the data are faceted by the levels of OC tendencies. These levels represent varying degrees of OCI-12 scores, approximating one standard deviation below the mean (Lower OC tendencies), at the mean (Medium OC tendencies), and one standard deviation above the mean (Higher OC tendencies), respectively. The color gradient used in the plot (yellow to pink to purple) visually reflects this increase in the level of OC tendencies. No relationship was observed between cognitive flexibility and RT-based statistical learning, and this pattern was consistent across the level of OC tendencies.

### How do OC tendencies modulate the interplay between cognitive flexibility and statistical learning? – Accuracy

We performed a linear mixed-effects model on blockwise mean accuracy of the ASRT task, with fixed effects for Time (Block 1-15), Triplet Type (high- vs. low-probability), OCI-12 score, Number of Perseverative errors of the Card Sorting Task, and all higher-order interactions. Random effects included participant-specific intercepts and correlated slopes for Time and Triplet Type to account for individual variability. The statistical parameters underlying the effects reported above are summarized in Table 2, whereas the full set of statistical estimates (b, SE, 95% CI, t, df, and p values) is provided in Supplementary Table S2.

**Table 2.**
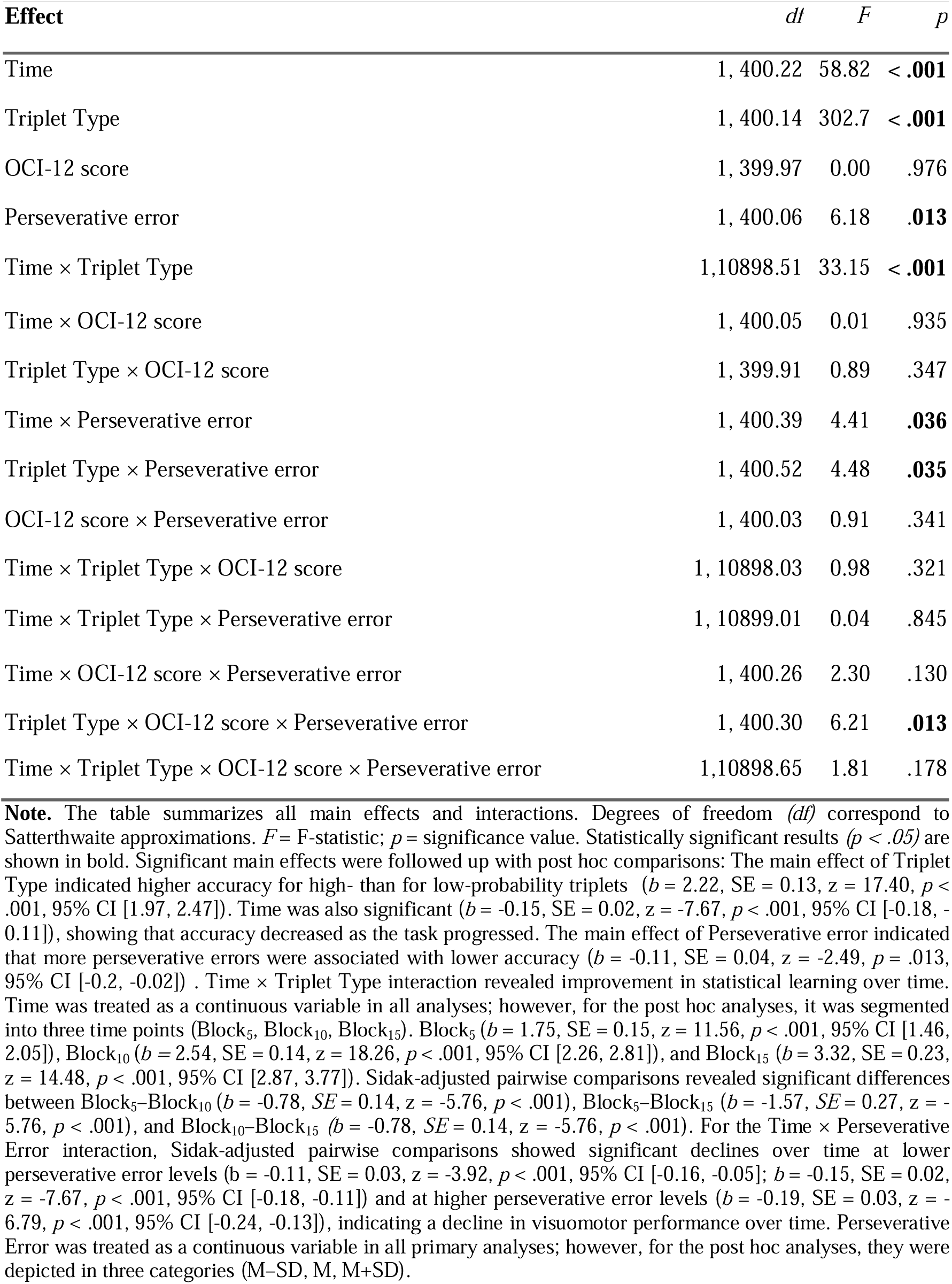
Results of the Linear Mixed-Effects Model predicting blockwise mean accuracy.

Participants were more accurate on high-probability trials than on low-probability trials, providing clear evidence of statistical learning (main effect of Triplet Type). Overall accuracy declined slightly over the course of the ASRT task (main effect of Time), but the difference between high- and low-probability trials increased, indicating a gradual improvement in statistical learning (Time × Triplet Type) on the ASRT task.

Cognitive flexibility influenced performance: a higher number of Perseverative errors on CST were significantly associated with lower overall accuracy (main effect of Perseverative error), and a modest decline in visuomotor performance over time on the ASRT task (Time × Perseverative error). Moreover, a significant Triplet Type × Perseverative error interaction revealed an antagonistic relationship between statistical learning and cognitive flexibility (Triplet Type × Perseverative error), with greater statistical learning linked to reduced cognitive flexibility, that is higher number of perseverative errors: (*b*_Low perseverative errors_ = 1.95, 95% CI [1.59, 2.30]; *b*_Mean perseverative errors_ = 2.22, 95% CI [1.97, 2.47]; b_High perseverative errors_ = 2.50, 95% CI [2.13, 2.86]). Pairwise comparisons indicated that the statistical learning increased significantly with perseverative errors (*b*_Low perseverative errors_ –_Mean perseverative errors_ = −0.28, 95% CI [−0.53, −0.02], *p* = .034; b_Low perseverative errors_ –_High perseverative errors_ = −0.55, 95% CI [−1.06, −0.04], *p* = .034; b_Mean perseverative errors_ –_High perseverative errors_ = −0.28, 95% CI [−0.53, −0.02], *p* = .034). Thus, individuals with poorer cognitive flexibility (higher numbers of perseverative errors) exhibited enhanced statistical learning, except when considering the trajectory of learning (Time × Triplet Type × Perseverative error), where this pattern was less pronounced.

OC tendencies did not significantly influence overall accuracy (main effect of OCI-12 score), visuomotor performance (Time × OCI-12 score), statistical learning (Triplet Type × OCI-12 score), or its improvement over time (Time × Triplet Type × OCI-12 score). Similarly, no interactions were observed between OC tendencies and Perseverative errors for overall accuracy or visuomotor performance (Time × Perseverative error × OCI-12 score). However, a significant Triplet Type × Perseverative error × OCI-12 score interaction indicated that OC tendencies modulate the antagonistic relationship between statistical learning and cognitive flexibility (see Figure 2). The post hoc analysis examining the trend of statistical learning across Perseverative error levels (M-SD, M, M+SD) revealed that among individuals with lower OC tendencies, Perseverative errors on CTS were positively associated with statistical learning, with a significant slope (*b* = 0.145, SE = 0.048, *z* = 2.97, 95% CI [0.050, 0.242], *p* = .002). A similar but weaker positive relationship was observed at the medium level of OC tendencies (*b* = 0.069, SE = 0.032, *z* = 2.11, 95% CI [0.005, 0.134], *p* = .034), suggesting that poorer cognitive flexibility was linked to enhanced statistical learning among lower and medium levels of OC tendencies. In contrast, this association disappeared among individuals with higher OC tendencies, where the slope was not statistically significant (*b* = −0.006, SE = 0.040, *z* = −0.17, 95% CI [−0.086, 0.072], *p* = .862). These coefficients indicate effects of modest magnitude and should be interpreted as reflecting relatively small differences between conditions. Overall, statistical learning increased as the number of perseverative errors rose for individuals with lower to medium OC tendencies, reflecting an antagonistic pattern in which reduced cognitive flexibility was accompanied by better statistical learning. This antagonistic pattern weakened in participants with high OC tendencies, where statistical learning remained moderate regardless of cognitive flexibility. Importantly, this modulation was not observed when examining the full trajectory of learning over time (Triplet Type × Time × Perseverative error × OCI-12 score).

**Figure 2.**
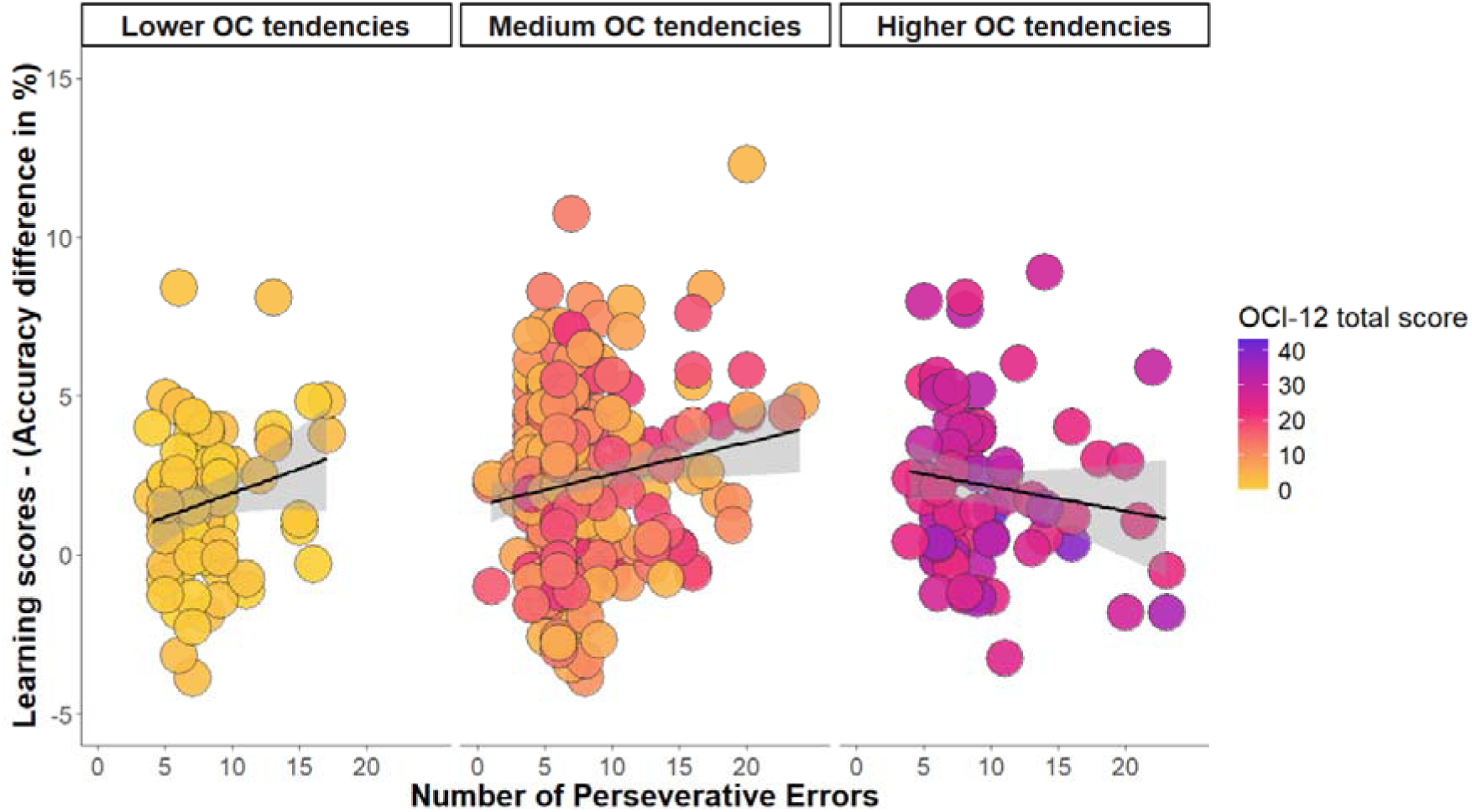
The figure illustrates the relationship between cognitive flexibility, as measured by the number of perseverative errors on the Card Sorting Task along the x-axis, and statistical learning performance along the y-axis, which is operationalized as the difference in accuracy between high- and low-probability triplets. Each point represents an individual participant, with color encoding their total OCI-12 score along a continuous gradient from low (yellow) through intermediate (pink) to high (purple) obsessive-compulsive tendencies. A linear regression line with 95% confidence intervals is overlaid in black. While the OCI-12 score was treated as a continuous variable in all statistical analyses, for visualization purposes, the data are faceted by the levels of OC tendencies. These levels represent varying degrees of OCI-12 scores, approximating one standard deviation below the mean (Lower OC tendencies) at the mean (Medium OC tendencies), and one standard deviation above the mean (Higher OC tendencies), respectively. The color gradient used in the plot (yellow to pink to purple) visually reflects this increase in the level of OC tendencies. The figure highlights the moderating role of OC tendencies in the antagonistic association between statistical learning and cognitive flexibility. Specifically, the typical inverse relationship, where reduced cognitive flexibility is associated with enhanced statistical learning, was observed in participants with Lower and Medium OC tendencies. In contrast, this pattern was not present among individuals with Higher OC tendencies, for whom statistical learning remained relatively stable across the cognitive flexibility continuum. This suggests a disruption of the typical trade-off between these cognitive functions in the context of OC tendencies.

## Discussion

In this study, we addressed a critical gap in understanding the relationship between goal-directed and automatic behaviour by examining how unsupervised SL interacts with cognitive flexibility across individuals. Rather than focusing on within-task dynamics, we tested whether this relationship generalizes across functions and is systematically modulated by OC tendencies. We conducted online experiment in a non-clinical university sample, assessing OC tendencies dimensionally across a continuous spectrum. SL was evaluated using an unsupervised learning task while cognitive flexibility was assessed with the Card Sorting Task. Our findings corroborate previous theories indicating a competitive interaction between SL and executive functions, specifically cognitive flexibility. Importantly, this relationship was not consistent across individuals: the inverse association between SL and cognitive flexibility weakened as OC tendencies increased. This pattern suggests that variability in OC tendencies fundamentally alters how automatic and controlled processes interact across tasks, rather than reflecting a simple shift in the dominance of one system over the other.

Our results build upon existing theories that highlight a disruption between automatic and goal-directed processes in both clinical OCD and OC tendencies or compulsivity ^8–12,24,28^ In particular, the typical antagonistic relationship between these functions, as emphasized by competitive memory system frameworks ^6,33,36–38,40,55–57^ appears to weaken with the increase of OC tendencies. This pattern may appear to challenge the habit hypothesis ^8,9,13,14,28^, which posits a shift toward habitual control under conditions of impaired goal-directed processing. However, methodological differences may partly account for these discrepancies since they predominantly rely on within-task paradigms, such as outcome devaluation or contingency degradation, which assess goal-directed behavior through adaptation to altered contingencies following extended instrumental training within a single context ^8,9,28^. Such paradigms often require the coordination of multiple processes simultaneously, and therefore capture how performance shifts under increased task demands. By contrast, the present findings point to a different form of alteration: the individual functions themselves remain largely intact, but their interaction changes with the increase of OC tendencies. Specifically, at lower levels of OC tendencies, SL and cognitive flexibility exhibit the expected antagonistic interplay, consistent with competitive memory frameworks ^6,36,58^. As OC tendencies increase, however, this relationship progressively diminishes, indicating that these processes no longer effectively constrain or inform one another. Importantly, this pattern does not reflect a simple shift in dominance, whereby one system overrides the other. Rather, it suggests a breakdown in their coordinated interaction, such that automatic and controlled processes operate more independently, without the typical antagonistic modulation. From this perspective, OC tendencies may be better characterized by altered coordination between cognitive systems than by dysfunction within any single function.

Taken together, the present results indicate that increasing OC tendencies are associated with a weakening of the relationship between executive control and SL. Importantly, the current design allowed for the independent assessment of these functions, enabling us to characterize not only their individual integrity but also their interaction. This distinction is critical for disentangling whether maladaptive behavioral patterns arise from enhanced automaticity^29,59,60^, reduced goal-directed control^11^, or alterations in the coordination between these systems ^61^. This perspective aligns with recent interaction-based accounts, which emphasize that cognitive inflexibility and habitual behaviour act as independent yet interacting predictors of OC symptomatology ^20^. Notably, prior work has demonstrated that these processes exert a compensatory interaction, such that both heightened habitual tendencies and reduced flexibility are jointly required for elevated symptom expression, while neither is sufficient in isolation^20^. At the same time, meta-analytic evidence challenges the assumption that executive or goal-directed functions are independently impaired in OCD, suggesting that observed deficits may reflect broader performance-related factors rather than specific functional dysfunctions^62^. In this context, our findings provide a complementary perspective: rather than reflecting deficits within individual systems, increasing OC tendencies are associated with a disruption in the functional relationship between them. Specifically, while SL and cognitive flexibility exhibit the expected antagonistic interplay at lower levels of OC tendencies, this relationship progressively diminishes as OC tendencies increase. These findings underscore the importance of moving beyond single-task paradigms and adopting cross-task approaches to capture how interactions between cognitive systems are organized across contexts. Future research employing multiple experimental paradigms will be essential to determine how such interaction-level alterations represent a core mechanism underlying OC or, in the long term, clinical OCD.

Our study identified the interaction between SL and the prefrontal cortex-related function, cognitive flexibility, with an across-task design. Nonetheless, the competitive memory systems hypothesis between executive functions and SL provides a theoretical framework that supports their interaction across four distinct levels^55^. At the most general level, correlational behavioral studies probe the relationship between executive functions and SL by employing independent tasks and testing for associations between their performance indices ^6,38,58,63^. In line with the present investigation, this approach targets enduring individual differences rather than transient states ^55^. Even at this level, it captures core neurocognitive strategies: when prefrontal, control-dependent processes are less efficient, SL systems appear to compensate by shifting toward more implicit, automatized modes of processing. The second level adopts an interventional approach, in which executive functions are transiently disrupted or downregulated to test how reduced top-down control influences SL performance ^36,37,64^. This design supports stronger causal inferences about their interaction. At the third level, converging neurocognitive evidence indicates that similar reductions in frontal control can emerge spontaneously, without external intervention, and may likewise facilitate statistical learning when control-related networks become less engaged ^65,66^. Finally, process-level approaches integrate executive functions and SL within a single task, enabling the simultaneous examination of their interaction under heightened cognitive demands ^55,56^. By combining the core features of both systems, such paradigms capture real-time, state-dependent dynamics as they unfold during performance. Collectively, evidence across these four levels converges on the conclusion that executive functions and SL interact in systematic ways, with their relationship flexibly shifting between competition and cooperation depending on contextual demands ^55^. Accordingly, to better understand how OC tendencies relate to this interplay, future research should aim to investigate all four levels within a unified empirical framework.

Beyond demonstrating an individual-difference-dependent interaction between neurocognitive systems, our findings address an important gap by elucidating the interaction between unsupervised learning, a primary form of learning ^30^, and executive functions among OC tendencies. Consistent with prior findings, unsupervised learning was found to be preserved ^21,67–70^, suggesting this core learning mechanism remains robust across OC tendencies similar to OCD, although not all prior findings are fully aligned with this pattern ^71–75^. This preservation is particularly noteworthy, as the excessive bias toward habitual behaviour in OCD may partly stem from external reward-driven learning. In feedback- or reward-driven contexts, it is difficult to disentangle whether performance improvements reflect sensitivity to external reinforcement or strengthened stimulus–response associations. Moreover, individuals with OCD and OC tendencies may reach a ceiling effect more rapidly due to heightened reward or feedback sensitivity, thereby accelerating the shift toward enhanced automatic processes and reducing the opportunity for flexible behavioral adaptation. Although the pure learning mechanism remained intact, the interaction of SL with goal-directed functions became atypical among higher OC tendencies. This suggests that the disruption lies not within the pure learning mechanisms themselves, but in the dynamic interplay between these systems among OC tendencies and possibly, in OCD ^9,28,61,76^. Future research should further investigate the within-task interaction between habitual and goal-directed processes in relation to unsupervised learning mechanisms, particularly in the absence of external reward. This would clarify the fundamental relationship between habitual and goal-directed functions and their contribution to OC- and OCD-related behavioral manifestations.

Our findings underscore the importance of investigating not only individual neurocognitive functions but also their dynamic interplay in advancing our understanding of the roots of OC tendencies. Our results imply that the weakened interplay observed among higher OC tendencies could serve as a potential predictor of OC symptomatology ^9,11,13,20,28^. Alongside theoretical frameworks that propose a bias toward automatic processes in clinical OCD ^9,10,13,20,27,28,76,77^, other perspectives emphasize shifts in the relative engagement of neurocognitive systems, whereby performance may be supported by increased reliance on controlled, explicit processes to maintain effective performance ^35,67,68,71,78,79^. Prior behavioral and neuroimaging studies have shown that during unsupervised tasks, neural activity is predominantly associated with hippocampal and medial temporal lobe regions rather than striatal structures ^69,70^. Such patterns reflect differential contributions of these systems during task performance. While the typical antagonistic interaction between these systems is maintained among lower levels of OC tendencies, this pattern appears to weaken as tendencies increase within the non-clinical OC spectrum. This raises the possibility that, in clinical populations, such a shift could result in an increased reliance on explicit systems ^67^. Our findings align with this perspective, as they suggest that the characteristics of both clinical OCD and OC tendencies may stem from the interactive manner of neurocognitive systems ^9,67,68,71,78,79^. Furthermore, it can be speculated that whether these systems engage in competitive or cooperative interactions depends on the specific cognitive demands of the task, suggesting a context-sensitive dynamic rather than a static relationship. Therefore, future research is needed to determine whether this altered cognitive interplay persists in clinical OCD or whether clinical OCD involves different interaction dynamics between these cognitive processes.

Another noteworthy finding is that the unrevealed interplay with cognitive flexibility appeared only in the accuracy-based performance indicator of the ASRT task. This dissociation is particularly interesting as it raises the question of which performance components are more sensitive to OC tendencies. One possible explanation is that the Card Sorting Task relies on accuracy without imposing temporal constraints ^46^. Accuracy- and rule-based learning generally require more effortful processing ^80–82^, which may account for the antagonistic relationship between the two measures. Specifically, enhanced implicit adaptation to rule-based structures may come at the cost of reduced performance in tasks requiring explicit rule detection, an effect that aligns with the competition and cooperation framework outlined above ^6,33,37,38,56,57^. This pattern also reflects the clinical features of OCD, where symptoms often involve an excessive focus on order, regularity, and precision, such as checking, arranging, or striving for symmetry ^7,83^. Consequently, accuracy-based measures may be particularly sensitive to the influence of OC tendencies, rather than reaction time. In our analyses using the Obsessive-Compulsive Inventory-Revised, individuals with higher OC scores exhibited faster reaction times. However, this effect did not replicate when using the OCI-12, a shortened version aligned with the DSM-5 ^7,54,84,85^. Given that the OCI-12 is derived from the same item pool, the disappearance of the effect following the removal of a subset of items suggests that the other finding with the OCI-R was not robust and is sensitive to the operationalization of the OC measure. Taken together, these findings suggest that OC symptoms may interfere more strongly with learning processes grounded in precision and rule detection, offering a clearer picture of how cognitive rigidity manifests in the disorder.

It is important to note that our study was conducted on an analogue sample, the so-called non-clinical adult population. Although OC tendencies in the general population share important phenomenological features with clinical OCD, and thus provide a relevant basis for theoretical inference, the current findings should not be generalized to clinical populations. Rather, they are best interpreted within a continuum-based framework. Spectrum approaches offer valuable theoretical and methodological tools for understanding clinical conditions, including OCD, by conceptualizing symptoms as continuously distributed and partially overlapping across individuals ^11,15,21,25,86,87^. Importantly, previous research has demonstrated that OC tendencies can influence a broad range of neurocognitive functions, including statistical learning ^21^, inhibitory control ^88^, cognitive flexibility^20^, and decision-making ^89,90^. This can prove that this approach serves as an effective methodological tool for identifying individual differences in neurocognitive functions. The current study contributes to this growing body of literature by highlighting how OC tendencies may influence not just cognitive domains in isolation but the dynamic interplay between them, offering a more integrated understanding of the neurocognitive profile associated with OC tendencies. Our findings indicate that while a dynamic interplay among neurocognitive functions is maintained at lower levels of OC tendencies, this interaction may potentially be absent in the clinical populations, resulting in impaired adaptive behavior. This disruption could be anticipated by higher OC tendencies. It is also possible that this inverse relationship between these functions persists at the clinical level but manifests in an exaggerated, potentially nonlinear manner, consistent with a U-shaped function across the OC spectrum.

In this study, OC tendencies were assessed using a multidimensional scale comprising washing, checking, ordering, and obsessing, with analyses focusing on the variance shared across these domains. While this approach captures core OC-related variation, it does not allow firm conclusions regarding the specificity of the observed associations. In particular, it remains to be determined to what extent these patterns reflect processes specific to OC symptoms, as opposed to broader transdiagnostic factors such as anxiety or intrusive thought propensity. More broadly, these findings are consistent with contemporary dimensional frameworks of psychopathology, including the Research Domain Criteria (RDoC) ^26,91–93^ and the Hierarchical Taxonomy of Psychopathology (HiTOP) ^94–96^, which emphasize that psychiatric traits are continuously distributed and organized along interacting neurocognitive systems. Future work incorporating multiple symptom dimensions will be important for clarifying the specificity and generality of the observed effects.

Some limitations should be noted in the present study. First, the relatively high exclusion rate, a common challenge in online experimental designs ^21,56,66,97–99^, may have affected the generalizability of the findings. Although rigorous data quality checks were essential to obtain high-quality data, the inclusion criterion requiring the absence of any self-reported psychiatric or neurological history likely limited the representation of individuals with more pronounced OC tendencies. As a result, the upper range of the OC spectrum may have been underrepresented, potentially narrowing the scope of our conclusions. A key limitation of the present study is that we did not control for other conditions that are frequently associated with elevated compulsivity and are often comorbid with OC tendencies, including anxiety and depressive symptomatology ^11,100,101^. Consequently, the present results do not allow us to fully isolate effects specifically related to the OC symptom dimension from those potentially driven by related affective or psychopathological factors. An additional limitation is that some participants exceeded commonly used clinical cutoffs on OC measures; however, in the absence of diagnostic assessment, it remains unclear whether these individuals meet the criteria for a formal OCD diagnosis. Presenting the tasks in a fixed order may have introduced potential order or fatigue effects. Future studies should therefore consider counterbalancing task order to minimize such influences, as performance on later tasks may be affected by declining attention or cognitive fatigue. ^102^ Moreover, although cognitive flexibility was assessed using a card sorting task, this measure has inherent methodological limitations ^102,103^. Therefore, future research would benefit from incorporating multiple measures of cognitive flexibility to obtain a more comprehensive assessment of executive functioning. Finally, the present study was conducted in a university-based adult sample, which may limit the generalizability of the findings to the broader population. Accordingly, future studies should extend these investigations to more diverse age groups and community samples to strengthen external validity.

Furthermore, OC tendencies were assessed using the OCI-based questionnaire, a widely used and psychometrically well-established measure. However, the OCI-R and its short forms consist of unidirectional items and do not include reverse-coded items. This feature has recently been identified as a potential source of bias in online studies. Sarna et al.^104^ argue that questionnaires without reverse-coded items are more susceptible to acquiescence, that is, a general tendency to agree with items regardless of their content. In addition, they show that inattentive responding can inflate the endorsement of low base rate symptoms, which may lead to artificially elevated symptom scores and, in turn, to spurious associations with behavioral measures. Related concerns have also been raised by Zorowitz et al. ^105^, who demonstrate that in online samples, careless or low effort responding can induce spurious correlations between task performance and psychiatric symptom measures, particularly when symptom distributions are positively skewed. Importantly, their findings suggest that standard quality control procedures based solely on behavioral task performance are often insufficient, as participants who respond inattentively on questionnaires are not reliably identified by behavioral exclusion criteria ^105^. In the present study, we applied rigorous behavioral screening procedures to ensure high-quality task data. At the same time, we did not include questionnaire-level attention checks or infrequency items within the OCI measure. The only potential proxy for estimating such effects is the Autism Quotient ^106^, as no additional questionnaires containing reverse-coded items were administered that would have allowed for the computation of an acquiescence index. Consequently, we cannot fully rule out the influence of acquiescence or inattentive responding on the observed associations. It is important to note that this limitation reflects a structural characteristic of the OCI instrument rather than a design choice specific to the present study. Addressing it more fully would likely require the development and validation of revised OC symptom measures that incorporate reverse-coded items while maintaining strong psychometric properties. Future studies should therefore combine well- established clinical instruments with improved questionnaire design features, including reverse-coded items and embedded attention checks, in order to more directly assess and control for response biases in online data collection contexts.

To conclude, our results support the competitive memory systems hypothesis, that emphasize an antagonistic relationship between neurocognitive systems ^6,33,36,40,57^ by elucidating a more general, individual-difference-dependent dynamic interplay between SL and goal-directed processes, here cognitive flexibility, and how their interaction is modulated by the prevalence of OC tendencies in a non-clinical population. While lower prevalence of OC tendencies may facilitate adaptive neurocognitive interactions, higher OC tendencies might represent early neurocognitive markers predictive of increased vulnerability to OCD. These results underscore the significance of conceptualizing OC tendencies as transdiagnostic traits to enhance early identification and inform tailored interventions. Importantly, our study highlights the necessity for cognitive neuroscience to transcend the examination of isolated neurocognitive functions and instead prioritize the investigation of their interactive dynamics, as the integrity of individual processes alone may mask critical dysfunctions emerging from their interactions.

## Resource availability Lead contact

Further information and requests for resources should be directed to and will be fulfilled by Prof. Dezső Németh.

## Materials availability

This study did not generate new unique reagents.

## Data availability statement

Data and codes for analysis are available on the following link: https://osf.io/fwb38/?view_only=03b2754e647742a2a8e5158a39174b45

## Acknowledgement

This work was supported by the the French National Grant Agency (ANR-24-CE37-5807) (to D.N.); the National Brain Research Program project NAP2022-I-2/2022 (to D.N.), EKÖP-24-3-II-ELTE-1159 (to B.B); the Spanish Ministry of Science, Innovation and Universities (MICIU), the State Research Agency (AEI), and the European Regional Development Fund (FEDER, UE) through the grant PID2024-160183NA-I00 (MICIU/AEI/10.13039/501100011033/FEDER, UE) (to T.V.)

## Contributions

B. B.: Conceptualization, Software, Validation, Investigation, Data Curation, Formal analysis, Writing – Original Draft, Writing – Review & Editing, Visualization; T.V.: Conceptualization, Software, Validation, Investigation, Data Curation, Formal analysis, Resources, Writing – Review & Editing, Supervision; B. C. F.: Methodology, Validation, Writing – Review & Editing, F. H.: Writing – Review & Editing; D.N.: Conceptualization, Software, Validation, Resources, Writing – Review & Editing, Supervision, Funding.

## Declaration of generative AI and AI-assisted technologies in the writing process

During the preparation of this work the author(s) used ChatGPT 4.0 to improve the writing style of our proposal. The author(s) reject using AI for scientific purposes to gain unethical benefits. However, the author(s) believe that it helps the equality of native and non-native English speakers to have the same opportunities. Thus, the author(s) do recommend using generative AI for lingual purposes (only). After using this tool/service, the author(s) reviewed and edited the content as needed and take(s) full responsibility for the content of the publication.

## STAR METHODS

### EXPERIMENTAL MODEL AND STUDY PARTICIPANT DETAILS

A total of 568 university students took part in the study through an online experiment in exchange for course credit. The sample shows partial participant overlap with those reported in Brezóczki et al. ^21^, Hann et al. ^87^, and Nagy et al. ^97^; however, the datasets are not identical, as data were collected across multiple studies with different primary research aims and measurement protocols. To maintain data integrity, rigorous quality control procedures were implemented, resulting in the exclusion of 164 participants who failed to follow the instructions. Our goal was to meet multiple data quality criteria, which are outlined in detail in the ‘Quality control of the data’ section below. The final sample consisted of 404 participants (285 female, 111 male, and 3 non-specified; *M_Age_ = 22.06 years; SD_Age_ = 5.09)*. This study did not perform analyses based on sex or gender, which could limit the generalizability of the findings. Participants provided informed consent, and the studies received approval from the Research Ethics Committee of Eötvös Loránd University, Budapest, Hungary (2021/504), following the principles of the Declaration of Helsinki.

### METHOD DETAILS

#### Alternating Serial Reaction Time task

Statistical learning was measured by the Alternating Serial Reaction Time (ASRT) task ^43,44^. The task was implemented in JavaScript using jsPsych version 6.1.0 ^107^ . Each trial presented a visual stimulus, an illustration of a dog’s head, in one of four horizontally aligned screen positions (Figure 3A). Participants were instructed to respond as quickly and accurately as possible by pressing the corresponding keyboard key (“S,” “F,” “J,” or “L” from left to right). The stimulus remained visible until a correct response was given, followed by a 120 ms response-to-stimulus interval before the next trial. Incorrect responses resulted in the stimulus remaining on the screen until corrected. Unbeknownst to participants, the stimuli followed a probabilistic eight-element sequence alternating between pattern and random elements (e.g., 2–r–4–r–3–r–1–r, where numbers denote fixed positions and “r” indicates a random one) (Figure 3B). This alternating structure resulted in some three-element combinations (triplets) occurring more frequently than others. High-probability triplets – such as 2–X–4, 4–X–3, 3–X–1, and 1–X–2, where X is the variable middle element – occurred more often due to their consistent alignment with the underlying sequence. High-probability triplets emerged either when two pattern elements enclosed a random element (50% of trials) or when two random elements enclosed a pattern element (12.5% of trials) (Figure 3C). Making up the remaining 37.5% of trials, low-probability triplets included less predictable element combinations based on the sequence structure.

**Figure 3.**
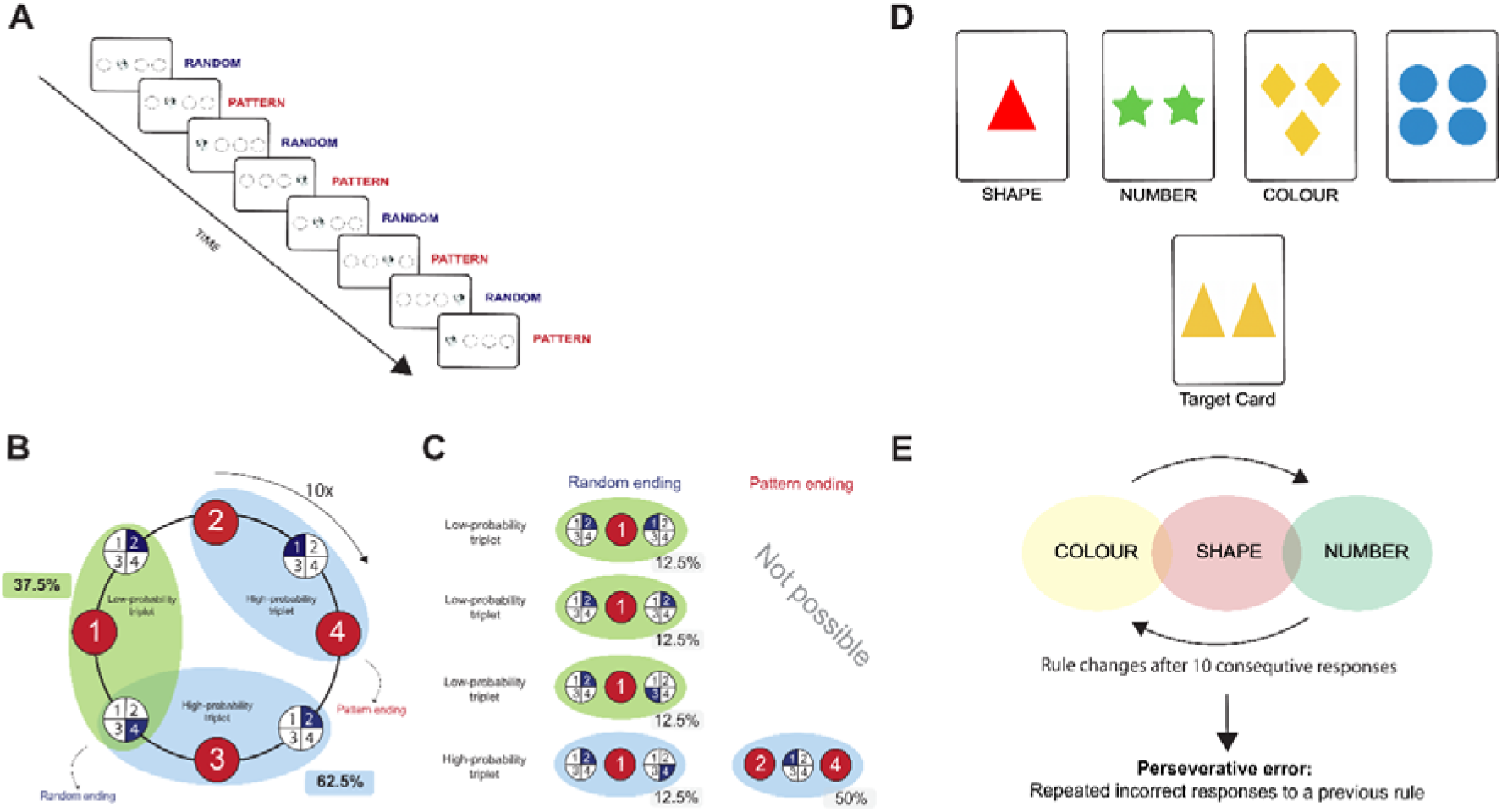
Design of the Study. **(A)** In the Alternating Serial Reaction Time (ASRT) task, participants were instructed to respond by pressing keys corresponding to the spatial location of a target stimulus, depicted as a dog’s head. Each trial followed an eight-element probabilistic sequence composed of alternating pattern and random elements. For example, a sequence such as 2–r–4–r–3–r–1–r consisted of fixed pattern trials at positions 2, 4, 3, and 1 (from left to right), interleaved with random elements, denoted by ‘r’, which could appear at any of the four possible locations. **(B)** Each trial was considered the third element in a sequence of three consecutive trials (a triplet). Pattern elements, which consistently appeared in fixed positions throughout the task, were displayed with red backgrounds, whereas random elements, selected randomly from the four possible locations, were shown with blue backgrounds. The structure of the sequence resulted in certain triplets occurring more frequently than others. These were categorized as high-probability triplets, while less frequent combinations were defined as low-probability triplets. **(C)** For illustrative purposes, three triplets are highlighted: 2(P)–1(R)–4(P), a pattern-ending high-probability triplet; 2(R)–3(P)–4(R), a random-ending high-probability triplet; and 4(R)–1(P)–2(R), a random-ending low-probability triplet. High-probability triplets could occur in one of two ways: either with two pattern trials and a random trial in the middle (accounting for 50% of all trials), or with two random trials and a pattern trial in the middle (accounting for 12.5% of trials). Overall, 62.5% of trials constituted the final element of a high-probability triplet, whereas the remaining 37.5% served as the final element of a low-probability triplet. **(D)** Schematic illustration of the Card Sorting Task (CST). Four reference cards are presented simultaneously, differing along three stimulus dimensions (color, shape, and quantity). On each trial, a target card appears below the reference cards, and participants are instructed to match it to one of the reference cards according to an sorting rule. **(E)** The rules are presented in a fixed, repeating order for up to six categories: Color, Shape, Number. The sorting rule changes after ten consecutive correct responses. Cognitive flexibility is indexed by the number of perseverative errors, defined as continued application of a previously correct but currently invalid rule.

The ASRT task consisted of 15 blocks, each containing 10 repetitions of an eight-element sequence, for a total of 80 stimuli per block. After each block, participants received feedback on their average accuracy and reaction time, but not on their statistical learning performance. Still, no information was provided about their statistical learning performance, as they were unaware that the task involved detecting a hidden regularity.

#### Card Sorting Task

Cognitive flexibility was assessed using the Card Sorting Task (CST), adapted from the Wisconsin Card Sorting Test (WCST), a well-established measure of executive function ^46,108,109^ The CST evaluates an individual’s ability to shift cognitive strategies in response to changing task demands. During the task, four reference cards are displayed simultaneously on the screen. Each card contains visual stimuli that differ along three dimensions: color (red, yellow, green, or blue), quantity (one to four items), and shape (triangle, star, diamond, or circle). In each trial, a separate target card appears beneath the reference cards (Figure 3D). Participants are instructed to match the target card with one of the four reference cards according to a common feature, either color, quantity, or shape, by selecting the corresponding card. Importantly, the correct matching rule is not disclosed to the participant. Instead, participants are required to infer the rule based on feedback provided after each response, which indicates whether the selected card was correct or incorrect (Figure 3E). The task comprises a total of sixty-four trials. The underlying categorization rule changes after every sequence of ten consecutive correct responses. The order of rule changes follows a fixed sequence: first based on color, then shape, followed by quantity, and then repeats in the same order. Cognitive flexibility is measured by the number of perseverative errors, defined as repeated use of a previously correct but no longer valid rule. A lower number of perseverative errors indicates greater cognitive flexibility.

#### Obsessive-Compulsive Inventory

Obsessive-compulsive tendencies were assessed using the Obsessive-Compulsive Inventory–Revised (OCI-R, ^54,110^), a self-report questionnaire measuring the severity of symptoms on a 5-point Likert scale ranging from 0 (not at all) to 4 (extremely). Participants reported how much each symptom had bothered them during the previous month. For the present analyses, a 12-item version (OCI-12) was computed from the OCI-R by excluding the hoarding and neutralizing items, in line with the DSM-5 reclassification ^54^. The OCI-12 has demonstrated strong psychometric validity, including high internal consistency and test-retest reliability ^54^. Although it represents a relatively recent modification, its Hungarian version, derived from the validated Hungarian OCI-R, has not yet been independently validated. The Cronbach’s alpha in our sample indicated good internal consistency (α = 0.89). The total score can vary from 0 to 48. In our final sample, the observed scores ranged from 0 to 43 (M = 12.58; SD = 9.02); see Supplementary Figure S1. For transparency, the full OCI-R analyses are reported in the Supplementary Materials (Supplementary Table S7-S10).

### Procedures

The experiment was conducted using the Gorilla Experiment Builder platform (https://www.gorilla.sc) ^111^. It began with participants reading an informational briefing and providing informed consent. Subsequently, participants completed the ASRT task, which started with two practice blocks followed by 15 task blocks. The practice blocks consisted solely of random trials and were excluded from analysis. To assess participants’ awareness of any patterns, we administered post-task inquiries asking them to report any regularities they may have noticed during the ASRT task. Specifically, participants were questioned about their detection of hidden sequences or patterns and asked to elaborate if applicable. None were able to accurately describe the alternating sequence, indicating that the task remained implicit, consistent with prior research ^33,112^. Following the ASRT task, cognitive flexibility was evaluated using the CST, and obsessive-compulsive tendencies were measured with the OCI-12. Additionally, Autism Quotient for autistic traits ^106^, demographic data on socioeconomic status, sleep, health, and handedness were collected. To account for potential confounding variables associated with the online setting, participants provided concise details about their testing environment, which are thoroughly outlined in the ‘Quality Control of the Data’ section.

### Quality control of the data

Online data collection is known to present several methodological challenges, as outcomes may be influenced not only by the psychological constructs of interest but also by variability in participants’ response styles, engagement levels, and task completion strategies ^104^. To address these concerns, we implemented data quality control procedures following established recommendations for online experimental research ^99^. From the initial sample, we excluded participants who: 1) reported prior completion of the ASRT task (N = 6, 1.06%), 2) restarted or quit the experiment at any point (N = 28, 4.93%); 3) failed the attention test screening for non-compliance (N = 26, 4.40%); 4) performed their mean raw accuracy on the ASRT task was below 80%, as such low performance was considered indicative of insufficient task engagement rather than genuine performance (N = 24, 4.23%); 5) were over 55 years due to the potential influence of age-related cognitive decline on neurocognitive performance measures (N = 1, 0.17%), 6) reported neurological impairment or head injury, or reported a diagnosis of any psychiatric condition (N = 58, 10.21%); 7) reported having consumed alcohol within 6 hours, recreational drugs or psychoactive medications within 24 hours before the experiment (N = 38, 6.69%); 8) took long self-paced breaks between ASRT blocks, as such extended interruptions may indicate reduced task engagement (N = 4, 0.7%), 9) random button pressing during ASRT task (N = 3, 0.53%) and 10) participants who scored in the zero category on the CST, indicating random clicking during the task or incomplete understanding of it (N = 19, 3.34%). As some participants met multiple exclusion criteria, the combined criteria resulted in the exclusion of a total of 164 participants (28.87%).

### QUANTIFICATION AND STATISTICAL ANALYSIS

#### Data preparation and analysis

Data preprocessing, statistical analysis, and visualization were carried out in R version 4.4.2 ^113^ For the ASRT task, each trial was categorized according to whether it represented the final element of a high- or low-probability triplet. Only the first response for each trial was retained; subsequent responses were excluded to ensure response validity. Trials with reaction times (RTs) below 150 ms or above 1000 ms, as well as those falling beyond three absolute median deviations from each participant’s median RT, were removed to control for outliers (Brezóczki et al., 2025). Repetition trials (e.g., 2–2–2, 1–1–1) and trills (e.g., 2–1–2, 3–2–3) were excluded due to their potential to elicit prepotent motor responses unrelated to statistical learning processes ^33,114^. The first two trials of each block were also removed, as they did not complete a triplet structure. Furthermore, incorrect responses were excluded from RT analyses. Following these procedures, 17.54% of trials were excluded from accuracy analyses and 24.61% from RT analyses. These exclusion rates are consistent with those reported in online ASRT studies ^21,56,66,97,98^.

We utilized Linear Mixed Models (LMMs) for analyzing RT and accuracy performance. LMMs were fit with the mixed function from the afex package ^115^.Regarding statistical learning and visuomotor performance, we computed the median reaction times and mean accuracy of the ASRT task for each participant in every block. These measures were used as outcome variables in linear mixed models with the Time (Block 1–15), Triplet Type (last element of a high- vs. low-probability triplet), Cognitive flexibility (Perseverative error), OC tendencies (OCI-12 score), and their higher-order interactions as fixed effects in the model. The Time, OCI-12 score, and Perseverative error factors were treated as mean-centered continuous variables. The models also included participant-specific intercepts and correlated slopes for the within-subject variables of Time and Triplet Type, treated as random effects. For RTs, the final model achieved convergence using the participants as random intercepts with by-participant correlated slopes for the Block. For accuracy, the final model achieved convergence using the participants as random intercepts with by-participant correlated slopes for Triplet Type and Time. Additional analyses conducted on further variables (Trials to Complete First Category, Total Number of Correct Responses) are reported in the Supplementary Information (Tables S5–S8).

Post hoc analyses involved the computation of estimated marginal means and marginal trends for simple effects and simple slopes, utilizing the emmeans package ^116^. Significant interactions involving continuous predictors, specifically the OCI-12 score and Perseverative errors, were probed at three levels: the mean, as well as at ±1 standard deviation from their means. For the Time factor, probing was performed at the end of the 5th, 10th, and 15th blocks. Figures were created with the ggplot2.

In reporting the results, the main effects of Time are interpreted as improvement in visuomotor performance, both on RT and accuracy, regardless of probabilistic patterns. Meanwhile, the main effect of Triplet Type is seen as an indicator of the amount of implicit probabilistic learning, and the interaction between Time and Triplet Type represents the temporal trajectory of statistical learning throughout the task. Please note that, although we used the OCI-12 score as a continuous variable for all our analyses, for more illustrative data visualization and clarified post hoc analysis, we depicted results at lower OC tendencies (Mean OCI-12 score - SD), medium OC tendencies (Mean OCI-12 score) and higher OC tendencies (Mean OCI-12 score + SD), based on standard deviations from the mean. For the more complex interactions, the same approach was applied to the Number of Perseverative errors.

Hence, our models were structured according to the following general form in *afex* syntax:

Reaction time:

*MedianRT ∼ Time * Triplet Type * OCI-12 score * Perseverative error + (Time* | *Participant)*

Accuracy:

*MeanAccuracy ∼ Time * Triplet Type * OCI-12 score * Perseverative error + (Time + Triplet Type* | *Participant)*

## References

1. Diamond, A. (2013). Executive Functions. Annu. Rev. Psychol. 64, 135–168. 10.1146/annurev-psych-113011-143750.

2. Friedman, N.P., and Robbins, T.W. (2022). The role of prefrontal cortex in cognitive control and executive function. Neuropsychopharmacology 47, 72–89. 10.1038/s41386-021-01132-0.

3. Miyake, A., Friedman, N.P., Emerson, M.J., Witzki, A.H., Howerter, A., and Wager, T.D. (2000). The Unity and Diversity of Executive Functions and Their Contributions to Complex “Frontal Lobe” Tasks: A Latent Variable Analysis. Cogn. Psychol. 41, 49–100. 10.1006/cogp.1999.0734.

4. Ashby, F.G., Turner, B.O., and Horvitz, J.C. (2010). Cortical and basal ganglia contributions to habit learning and automaticity. Trends Cogn. Sci. 14, 208–215. 10.1016/j.tics.2010.02.001.

5. Kaufman, S.B., DeYoung, C.G., Gray, J.R., Jiménez, L., Brown, J., and Mackintosh, N. (2010). Implicit learning as an ability. Cognition 116, 321–340. 10.1016/j.cognition.2010.05.011.

6. Pedraza, F., Farkas, B.C., Vékony, T., Haesebaert, F., Phelipon, R., Mihalecz, I., Janacsek, K., Anders, R., Tillmann, B., Plancher, G., et al. (2024). Evidence for a competitive relationship between executive functions and statistical learning. NPJ Sci. Learn. 9, 30. 10.1038/s41539-024-00243-9.

7. American Psychiatric Association (2022). Diagnostic and Statistical Manual of Mental Disorders (American Psychiatric Association Publishing) 10.1176/appi.books.9780890425787.

8. Gillan, C.M., Apergis-Schoute, A.M., Morein-Zamir, S., Urcelay, G.P., Sule, A., Fineberg, N.A., Sahakian, B.J., and Robbins, T.W. (2015). Functional Neuroimaging of Avoidance Habits in Obsessive-Compulsive Disorder. American Journal of Psychiatry 172, 284–293. 10.1176/appi.ajp.2014.14040525.

9. Gillan, C.M., Papmeyer, M., Morein-Zamir, S., Sahakian, B.J., Fineberg, N.A., Robbins, T.W., and de Wit, S. (2011). Disruption in the Balance Between Goal-Directed Behavior and Habit Learning in Obsessive-Compulsive Disorder. American Journal of Psychiatry 168, 718–726. 10.1176/appi.ajp.2011.10071062.

10. Gillan, C.M., and Robbins, T.W. (2014). Goal-directed learning and obsessive–compulsive disorder. Philosophical Transactions of the Royal Society B: Biological Sciences 369, 20130475. 10.1098/rstb.2013.0475.

11. Gillan, C.M., Kosinski, M., Whelan, R., Phelps, E.A., and Daw, N.D. (2016). Characterizing a psychiatric symptom dimension related to deficits in goal-directed control. Elife 5. 10.7554/eLife.11305.

12. Yu, Q., Gao, F., Li, C., Xia, J., Cao, Y., Wang, X., Xiao, C., Lu, J., Liu, Q., Fan, J., et al. (2024). Compulsion is associated with impaired goal-directed and habitual learning and responding in obsessive-compulsive disorder. International Journal of Clinical and Health Psychology 24, 100531. 10.1016/j.ijchp.2024.100531.

13. Gillan, C.M. (2021). Recent developments in the habit hypothesis of OCD and compulsive disorders. The neurobiology and treatment of OCD: accelerating progress, 147–167.

14. Banca, P., Voon, V., Vestergaard, M.D., Philipiak, G., Almeida, I., Pocinho, F., Relvas, J., and Castelo-Branco, M. (2015). Imbalance in habitual versus goal directed neural systems during symptom provocation in obsessive-compulsive disorder. Brain 138, 798–811. 10.1093/brain/awu379.

15. Abramowitz, J.S., Fabricant, L.E., Taylor, S., Deacon, B.J., McKay, D., and Storch, E.A. (2014). The relevance of analogue studies for understanding obsessions and compulsions. Clin. Psychol. Rev. 34, 206–217. 10.1016/j.cpr.2014.01.004.

16. Hajcak, G., Huppert, J.D., Simons, R.F., and Foa, E.B. (2004). Psychometric properties of the OCI-R in a college sample. Behaviour Research and Therapy 42, 115–123. 10.1016/j.brat.2003.08.002.

17. Foa, E.B., Huppert, J.D., Leiberg, S., Langner, R., Kichic, R., Hajcak, G., and Salkovskis, P.M. (2002). The Obsessive-Compulsive Inventory: Development and validation of a short version. Psychol. Assess. 14, 485–496. 10.1037/1040-3590.14.4.485.

18. Fullana, M.A., Vilagut, G., Rojas-Farreras, S., Mataix-Cols, D., de Graaf, R., Demyttenaere, K., Haro, J.M., de Girolamo, G., Lépine, J.P., Matschinger, H., et al. (2010). Obsessive–compulsive symptom dimensions in the general population: Results from an epidemiological study in six European countries. J. Affect. Disord. 124, 291–299. 10.1016/j.jad.2009.11.020.

19. Mataix-Cols, D., Junqué, C., Sànchez-Turet, M., Vallejo, J., Verger, K., and Barrios, M. (1999). Neuropsychological functioning in a subclinical obsessive-compulsive sample. Biol. Psychiatry 45, 898–904. 10.1016/S0006-3223(98)00260-1.

20. Ramakrishnan, S., Robbins, T.W., and Zmigrod, L. (2022). Cognitive Rigidity, Habitual Tendencies, and Obsessive-Compulsive Symptoms: Individual Differences and Compensatory Interactions. Front. Psychiatry 13. 10.3389/fpsyt.2022.865896.

21. Brezóczki, B., Farkas, B.C., Hann, F., Pesthy, O., Tóth-Fáber, E., Farkas, K., Csigó, K., Németh, D., and Vékony, T. (2025). Individual differences in probabilistic learning and updating predictive representations in individuals with obsessive-compulsive tendencies. BMC Psychiatry 25, 368. 10.1186/s12888-025-06786-4.

22. Zermatten, A., Van der Linden, M., Jermann, F., and Ceschi, G. (2006). Validation of a French version of the Obsessive–Compulsive Inventory-Revised in a non-clinical sample. European Review of Applied Psychology 56, 151–155. 10.1016/j.erap.2005.07.003.

23. Rachman, S., and de Silva, P. (1978). Abnormal and normal obsessions. Behaviour research and therapy 16, 233–248.

24. Gillan, C.M., Kalanthroff, E., Evans, M., Weingarden, H.M., Jacoby, R.J., Gershkovich, M., Snorrason, I., Campeas, R., Cervoni, C., Crimarco, N.C., et al. (2020). Comparison of the Association Between Goal-Directed Planning and Self-reported Compulsivity vs Obsessive-Compulsive Disorder Diagnosis. JAMA Psychiatry 77, 77. 10.1001/jamapsychiatry.2019.2998.

25. Gillan, C.M., and Rutledge, R.B. (2021). Smartphones and the Neuroscience of Mental Health. Annu. Rev. Neurosci. 44, 129–151. 10.1146/annurev-neuro-101220-014053.

26. Colzato, L., Zhang, H., Roessner, V., Beste, C., and Hommel, B. (2025). Non-bivalent psychopathology: Rethinking mental disorders through metacontrol. Neurosci. Biobehav. Rev. 176, 106297. 10.1016/j.neubiorev.2025.106297.

27. Graybiel, A.M., and Rauch, S.L. (2000). Toward a Neurobiology of Obsessive-Compulsive Disorder. Neuron 28, 343–347. 10.1016/S0896-6273(00)00113-6.

28. Gillan, C.M., Morein-Zamir, S., Urcelay, G.P., Sule, A., Voon, V., Apergis-Schoute, A.M., Fineberg, N.A., Sahakian, B.J., and Robbins, T.W. (2014). Enhanced Avoidance Habits in Obsessive-Compulsive Disorder. Biol. Psychiatry 75, 631–638. 10.1016/j.biopsych.2013.02.002.

29. Banca, P., Herrojo Ruiz, M., Gonzalez-Zalba, M.F., Biria, M., Marzuki, A.A., Piercy, T., Sule, A., Fineberg, N.A., and Robbins, T.W. (2024). Action sequence learning, habits, and automaticity in obsessive-compulsive disorder. Elife 12. 10.7554/eLife.87346.4.

30. Zhong, L., Baptista, S., Gattoni, R., Arnold, J., Flickinger, D., Stringer, C., and Pachitariu, M. (2025). Unsupervised pretraining in biological neural networks. Nature. 10.1038/s41586-025-09180-y.

31. Aslin, R.N. (2017). Statistical learning: a powerful mechanism that operates by mere exposure. WIREs Cognitive Science 8. 10.1002/wcs.1373.

32. Conway, C.M. (2020). How does the brain learn environmental structure? Ten core principles for understanding the neurocognitive mechanisms of statistical learning. Neurosci. Biobehav. Rev. 112, 279–299. 10.1016/j.neubiorev.2020.01.032.

33. Horváth, K., Nemeth, D., and Janacsek, K. (2022). Inhibitory control hinders habit change. Sci. Rep. 12, 8338. 10.1038/s41598-022-11971-6.

34. Theeuwes, J., Bogaerts, L., and van Moorselaar, D. (2022). What to expect where and when: how statistical learning drives visual selection. Trends Cogn. Sci. 26, 860–872. 10.1016/j.tics.2022.06.001.

35. Brezóczki, B., Vékony, T., Pesthy, O., Tóth-Fáber, E., Csigó, K., Farkas, K., and Nemeth, D. (2023). Unraveling sequence learning in obsessive–compulsive disorder. Curr. Opin. Behav. Sci. 54, 101326. 10.1016/j.cobeha.2023.101326.

36. Ambrus, G.G., Vékony, T., Janacsek, K., Trimborn, A.B.C., Kovács, G., and Nemeth, D. (2020). When less is more: Enhanced statistical learning of non-adjacent dependencies after disruption of bilateral DLPFC. J. Mem. Lang. 114, 104144. 10.1016/j.jml.2020.104144.

37. Nemeth, D., Janacsek, K., Polner, B., and Kovacs, Z.A. (2013). Boosting Human Learning by Hypnosis. Cerebral Cortex 23, 801–805. 10.1093/cercor/bhs068.

38. Virag, M., Janacsek, K., Horvath, A., Bujdoso, Z., Fabo, D., and Nemeth, D. (2015). Competition between frontal lobe functions and implicit sequence learning: evidence from the long-term effects of alcohol. Exp. Brain Res. 233, 2081–2089. 10.1007/s00221-015-4279-8.

39. Tóth-Fáber, E., Janacsek, K., Szőllősi, Á., Kéri, S., and Nemeth, D. (2021). Regularity detection under stress: Faster extraction of probability-based regularities. PLoS One 16, e0253123. 10.1371/journal.pone.0253123.

40. Poldrack, R.A., Clark, J., Paré-Blagoev, E.J., Shohamy, D., Creso Moyano, J., Myers, C., and Gluck, M.A. (2001). Interactive memory systems in the human brain. Nature 414, 546–550. 10.1038/35107080.

41. Chamberlain, S.R., Fineberg, N.A., Menzies, L.A., Blackwell, A.D., Bullmore, E.T., Robbins, T.W., and Sahakian, B.J. (2007). Impaired Cognitive Flexibility and Motor Inhibition in Unaffected First-Degree Relatives of Patients With Obsessive-Compulsive Disorder. American Journal of Psychiatry 164, 335–338. 10.1176/ajp.2007.164.2.335.

42. Tomiyama, H., Nakao, T., Murayama, K., Nemoto, K., Ikari, K., Yamada, S., Kuwano, M., Hasuzawa, S., Togao, O., Hiwatashi, A., et al. (2019). Dysfunction between dorsal caudate and salience network associated with impaired cognitive flexibility in obsessive-compulsive disorder: A resting-state fMRI study. Neuroimage Clin. 24, 102004. 10.1016/j.nicl.2019.102004.

43. Farkas, B.C., Krajcsi, A., Janacsek, K., and Nemeth, D. (2023). The complexity of measuring reliability in learning tasks: An illustration using the Alternating Serial Reaction Time Task. Behav. Res. Methods 56, 301–317. 10.3758/s13428-022-02038-5.

44. Howard, J.H., and Howard, D. V. (1997). Age differences in implicit learning of higher order dependencies in serial patterns. Psychol. Aging 12, 634–656. 10.1037/0882-7974.12.4.634.

45. Nemeth, D., and Tóth-Fáber, E. (2026). Resolving Fundamental Debates in Statistical Learning: The Power of Process Dissociation. Preprint, 10.31234/osf.io/uwa35_v1.

46. Fox, C.J., Mueller, S.T., Gray, H.M., Raber, J., and Piper, B.J. (2013). Evaluation of a Short-Form of the Berg Card Sorting Test. PLoS One 8, e63885. 10.1371/journal.pone.0063885.

47. Zhang, Z., Yang, L.-Z., Vékony, T., Wang, C., and Li, H. (2024). Split-half reliability estimates of an online card sorting task in a community sample of young and elderly adults. Behav. Res. Methods 56, 1039–1051.

48. Baker, K.S., Gibson, S.J., GeorgiouLKaristianis, N., and Giummarra, M.J. (2018). Relationship between selfLreported cognitive difficulties, objective neuropsychological test performance and psychological distress in chronic pain. European Journal of Pain 22, 601–613. 10.1002/ejp.1151.

49. Pesthy, Z.V., Berta, K., Vékony, T., Farkas, B.C., Németh, D., and Kun, B. (2026). Dissociating the cognitive underpinnings of recreational cannabis use from problematic use. Compr. Psychiatry 147, 152685. 10.1016/j.comppsych.2026.152685.

50. Berta, K., Pesthy, Z.V., Vékony, T., Farkas, B.C., Németh, D., and Kun, B. (2023). The neuropsychological profile of work addiction. Sci. Rep. 13, 20090. 10.1038/s41598-023-47515-9.

51. Wollenhaupt, C., Wilke, L., Erim, Y., Rauh, M., Steins-Loeber, S., and Paslakis, G. (2019). The association of leptin secretion with cognitive performance in patients with eating disorders. Psychiatry Res. 276, 269–277. 10.1016/j.psychres.2019.05.001.

52. Gelonch, O., Garolera, M., Valls, J., Rosselló, L., and Pifarré, J. (2016). Executive function in fibromyalgia: Comparing subjective and objective measures. Compr. Psychiatry 66, 113–122. 10.1016/j.comppsych.2016.01.002.

53. Dickson, K.S., Ciesla, J.A., and Zelic, K. (2017). The Role of Executive Functioning in Adolescent Rumination and Depression. Cognit. Ther. Res. 41, 62–72. 10.1007/s10608-016-9802-0.

54. Abramovitch, A., Abramowitz, J.S., and McKay, D. (2021). The OCI-12: A syndromally valid modification of the obsessive-compulsive inventory-revised. Psychiatry Res. 298, 113808. 10.1016/j.psychres.2021.113808.

55. Horváth, K., Nemeth, D., Janacsek, K., and Kóbor, A. (2025). Cooperative interaction between statistical learning and inhibitory control 10.31234/osf.io/ptdkr_v2.

56. Vékony, T., Brezóczki, B., Csifcsák, G., Németh, D., and Simor, P. (2025). A functional trade-off between executive control and implicit statistical learning is dynamically gated by mind wandering. Preprint, 10.1101/2025.08.05.668618.

57. Pedraza, F., Vékony, T., Farkas, B.C., Haesebaert, F., Phelipon, R., Mihalecz, I., Janacsek, K., Tillmann, B., Anders, R., Plancher, G., et al. (2024). The interplay between executive functions and updating predictive representations. Preprint, 10.1101/2024.12.05.626969.

58. Pedraza, F., Vékony, T., Farkas, B.C., Haesebaert, F., Phelipon, R., Mihalecz, I., Janacsek, K., Tillmann, B., Anders, R., Plancher, G., et al. (2025). The interplay between executive functions and updating predictive representations. Sci. Rep. 15, 30555. 10.1038/s41598-025-14876-2.

59. Barzilay, S., Fradkin, I., and Huppert, J.D. (2022). Habitual or hyper-controlled behavior: OCD symptoms and explicit sequence learning. J. Behav. Ther. Exp. Psychiatry 75, 101723. 10.1016/j.jbtep.2022.101723.

61. Donegan, K.R., Suddell, S., De Forest, S., Sun, J., Gallagher, E., Hanlon, A., Buabang, E.K., Conelea, C., Mackay, C., and Gillan, C.M. (2025). Compulsivity is associated with accelerated stimulus- response habit learning. Preprint, 10.31234/osf.io/xwtsu_v1.

61. Gruner, P., Anticevic, A., Lee, D., and Pittenger, C. (2016). Arbitration between Action Strategies in Obsessive-Compulsive Disorder. The Neuroscientist 22, 188–198. 10.1177/1073858414568317.

62. Fradkin, I., Strauss, A.Y., Pereg, M., and Huppert, J.D. (2018). Rigidly Applied Rules? Revisiting Inflexibility in Obsessive Compulsive Disorder Using Multilevel Meta-Analysis. Clinical Psychological Science 6, 481–505. 10.1177/2167702618756069.

63. Park, J., Yoon, H.-D., Yoo, T., Shin, M., and Jeon, H.-A. (2020). Potential and efficiency of statistical learning closely intertwined with individuals’ executive functions: a mathematical modeling study. Sci. Rep. 10, 18843.

64. Smalle, E.H.M., Daikoku, T., Szmalec, A., Duyck, W., and Möttönen, R. (2022). Unlocking adults’ implicit statistical learning by cognitive depletion. Proceedings of the National Academy of Sciences 119, e2026011119.

65. Simor, P., Vékony, T., Farkas, B.C., Szalárdy, O., Bogdány, T., Brezóczki, B., Csifcsák, G., and Németh, D. (2025). Mind Wandering during Implicit Learning Is Associated with Increased Periodic EEG Activity and Improved Extraction of Hidden Probabilistic Patterns. The Journal of Neuroscience 45, e1421242025. 10.1523/JNEUROSCI.1421-24.2025.

66. Vékony, T., Farkas, B.C., Brezóczki, B., Mittner, M., Csifcsák, G., Simor, P., and Németh, D. (2025). Mind wandering enhances statistical learning. iScience 28, 111703. 10.1016/j.isci.2024.111703.

67. Soref, A., Liberman, N., Abramovitch, A., Poznanski, Y., and Dar, R. (2021). Intact capacity for implicit learning in obsessive-compulsive disorder. J. Behav. Ther. Exp. Psychiatry 73, 101667. 10.1016/j.jbtep.2021.101667.

68. Soref, A., Liberman, N., Abramovitch, A., and Dar, R. (2018). Explicit instructions facilitate performance of OCD participants but impair performance of non-OCD participants on a serial reaction time task. J. Anxiety Disord. 55, 56–62. 10.1016/j.janxdis.2018.02.003.

69. Rauch, S.L., Wedig, M.M., Wright, C.I., Martis, B., McMullin, K.G., Shin, L.M., Cannistraro, P.A., and Wilhelm, S. (2007). Functional Magnetic Resonance Imaging Study of Regional Brain Activation During Implicit Sequence Learning in Obsessive–Compulsive Disorder. Biol. Psychiatry 61, 330–336. 10.1016/j.biopsych.2005.12.012.

70. Rauch, S.L., Savage, C.R., Alpert, N.M., Dougherty, D., Kendrick, A., Curran, T., Brown, H.D., Manzo, P., Fischman, A., and Jenike, M.A. (1997). Probing striatal function in obsessive-compulsive disorder: a PET study of implicit sequence learning. Journal of Neuropsychiatry and Clinical Neurosciences 9, 568–573.

71. Deckersbach, T., Savage, C.R., Curran, T., Bohne, A., Wilhelm, S., Baer, L., Jenike, M.A., and Rauch, S.L. (2002). A Study of Parallel Implicit and Explicit Information Processing in Patients With Obsessive-Compulsive Disorder. American Journal of Psychiatry 159, 1780–1782. 10.1176/appi.ajp.159.10.1780.

72. Kathmann, N., Rupertseder, C., Hauke, W., and Zaudig, M. (2005). Implicit Sequence Learning in Obsessive-Compulsive Disorder: Further Support for the Fronto-Striatal Dysfunction Model. Biol. Psychiatry 58, 239–244. 10.1016/j.biopsych.2005.03.045.

73. Goldman, B.L., Martin, E.D., Calamari, J.E., Woodard, J.L., Chik, H.M., Messina, M.G., Pontarelli, N.K., Marker, C.D., Riemann, B.C., and Wiegartz, P.S. (2008). Implicit learning, thought-focused attention and obsessive-compulsive disorder: A replication and extension. Behaviour Research and Therapy 46, 48–61. 10.1016/j.brat.2007.10.004.

74. Marker, C.D., Calamari, J.E., Woodard, J.L., and Riemann, B.C. (2006). Cognitive self-consciousness, implicit learning and obsessive–compulsive disorder. J. Anxiety Disord. 20, 389–407. 10.1016/j.janxdis.2005.03.003.

75. Vloet, T.D., Marx, I., Kahraman-Lanzerath, B., Zepf, F.D., Herpertz-Dahlmann, B., and Konrad, K. (2010). Neurocognitive Performance in Children with ADHD and OCD. J. Abnorm. Child Psychol. 38, 961–969. 10.1007/s10802-010-9422-1.

76. Robbins, T.W., Vaghi, M.M., and Banca, P. (2019). Obsessive-Compulsive Disorder: Puzzles and Prospects. Neuron 102, 27–47. 10.1016/j.neuron.2019.01.046.

77. Vaghi, M.M., Cardinal, R.N., Apergis-Schoute, A.M., Fineberg, N.A., Sule, A., and Robbins, T.W. (2019). Action-Outcome Knowledge Dissociates From Behavior in Obsessive-Compulsive Disorder Following Contingency Degradation. Biol. Psychiatry Cogn. Neurosci. Neuroimaging 4, 200–209. 10.1016/j.bpsc.2018.09.014.

78. Ullman, M.T. (2004). Contributions of memory circuits to language: the declarative/procedural model. Cognition 92, 231–270. 10.1016/j.cognition.2003.10.008.

79. Ullman, M.T., and Pullman, M.Y. (2015). A compensatory role for declarative memory in neurodevelopmental disorders. Neurosci. Biobehav. Rev. 51, 205–222.

80. Nomura, E., Maddox, W., Filoteo, J., Ing, A., Gitelman, D., Parrish, T., Mesulam, M.-M., and Reber, P. (2006). Neural Correlates of Rule-Based and Information-Integration Visual Category Learning. Cerebral Cortex 17, 37–43. 10.1093/cercor/bhj122.

81. Feng, X., Perceval, G.J., Feng, W., and Feng, C. (2020). High Cognitive Flexibility Learners Perform Better in Probabilistic Rule Learning. Front. Psychol. 11. 10.3389/fpsyg.2020.00415.

82. Seger, C.A., and Miller, E.K. (2010). Category Learning in the Brain. Annu. Rev. Neurosci. 33, 203–219. 10.1146/annurev.neuro.051508.135546.

83. Bragdon, L.B., Gibb, B.E., and Coles, M.E. (2018). Does neuropsychological performance in OCD relate to different symptoms? A meta-analysis comparing the symmetry and obsessing dimensions. Depress. Anxiety 35, 761–774. 10.1002/da.22785.

84. American Psychiatric Association., and American Psychiatric Association. DSM-5 Task Force. (2013). Diagnostic and statistical manual of mental disordersL: DSM-5. (American Psychiatric Association).

85. Leckman, J.F., Rauch, S.L., and Mataix-Cols, D. (2007). Symptom Dimensions in Obsessive-Compulsive Disorder: *Implications for the* DSM-V. CNS Spectr. 12, 376–387. 10.1017/S1092852900021179.

86. Gillan, C.M., Fineberg, N.A., and Robbins, T.W. (2017). A trans-diagnostic perspective on obsessive-compulsive disorder. Psychol. Med. 47, 1528–1548. 10.1017/S0033291716002786.

87. Hann, F., Pesthy, O., Brezóczki, B., Vékony, T., Nagy, C.A., Sapey-Triomphe, L.-A., Tóth-Fáber, E., Farkas, B.C., Farkas, K., and Németh, D. (2025). Autistic traits relate to speed/accuracy trade-off but not statistical learning and updating. Sci. Rep. 15, 32001. 10.1038/s41598-025-16138-7.

88. Dominke, C., Graham-Schmidt, K., Gentsch, A., and Schütz-Bosbach, S. (2021). Action inhibition in individuals with high obsessive-compulsive trait of incompleteness: An ERP study. Biol. Psychol. 159, 108019. 10.1016/j.biopsycho.2021.108019.

89. Scholl, J., Trier, H.A., Rushworth, M.F.S., and Kolling, N. (2022). The effect of apathy and compulsivity on planning and stopping in sequential decision-making. PLoS Biol. 20, e3001566. 10.1371/journal.pbio.3001566.

91. Hoven, M., Mulder, T., Denys, D., van Holst, R.J., and Luigjes, J. (2024). Lower confidence and increased error sensitivity in OCD patients while learning under volatility. Transl. Psychiatry 14, 370. 10.1038/s41398-024-03042-3.

91. Mittal, V.A., and Wakschlag, L.S. (2016). Research domain criteria (RDoC) grows up: Strengthening neurodevelopmental investigation within the RDoC framework. J. Affect. Disord. 216, 30.

92. Insel, T.R. (2014). The NIMH research domain criteria (RDoC) project: precision medicine for psychiatry. American journal of psychiatry 171, 395–397.

93. Insel, T., Cuthbert, B., Garvey, M., Heinssen, R., Pine, D.S., Quinn, K., Sanislow, C., and Wang, P. (2010). Research domain criteria (RDoC): toward a new classification framework for research on mental disorders. Preprint at American Psychiatric Association.

94. Kotov, R., Krueger, R.F., Watson, D., Achenbach, T.M., Althoff, R.R., Bagby, R.M., Brown, T.A., Carpenter, W.T., Caspi, A., and Clark, L.A. (2017). The Hierarchical Taxonomy of Psychopathology (HiTOP): A dimensional alternative to traditional nosologies. J. Abnorm. Psychol. 126, 454.

95. Kotov, R., Krueger, R.F., Watson, D., Cicero, D.C., Conway, C.C., DeYoung, C.G., Eaton, N.R., Forbes, M.K., Hallquist, M.N., and Latzman, R.D. (2021). The Hierarchical Taxonomy of Psychopathology (HiTOP): A quantitative nosology based on consensus of evidence. Annu. Rev. Clin. Psychol. 17, 83–108.

96. Kotov, R., Cicero, D.C., Conway, C.C., DeYoung, C.G., Dombrovski, A., Eaton, N.R., First, M.B., Forbes, M.K., Hyman, S.E., and Jonas, K.G. (2022). The Hierarchical Taxonomy of Psychopathology (HiTOP) in psychiatric practice and research. Psychol. Med. 52, 1666–1678.

97. Nagy, C.A., Hann, F., Brezóczki, B., Farkas, K., Vékony, T., Pesthy, O., and Németh, D. (2025). Intact ultrafast memory consolidation in adults with autism and neurotypicals with autism traits. Brain Res. 1847, 149299. 10.1016/j.brainres.2024.149299.

98. Hann, F., Pesthy, O., Brezóczki, B., Vékony, T., Nagy, C.A., Sapey-Triomphe, L.-A., Tóth-Fáber, E., Farkas, B.C., Farkas, K., and Nemeth, D. (2024). Autistic Traits Relate to Speed/Accuracy Trade-off but not Statistical Learning and Updating. Preprint, 10.31219/osf.io/p8m37_v1.

99. Rodd, J.M. (2024). Moving experimental psychology online: How to obtain high quality data when we can’t see our participants. J. Mem. Lang. 134, 104472. 10.1016/j.jml.2023.104472.

100. Choi, K.W., Kim, Y.-K., and Jeon, H.J. (2020). Comorbid Anxiety and Depression: Clinical and Conceptual Consideration and Transdiagnostic Treatment. In, pp. 219–235. 10.1007/978-981-32-9705-0_14.

101. Goodwin, G.M. (2015). The overlap between anxiety, depression, and obsessive-compulsive disorder. Dialogues Clin. Neurosci. 17, 249–260. 10.31887/DCNS.2015.17.3/ggoodwin.

102. Ord, J.S., Greve, K.W., Bianchini, K.J., and Aguerrevere, L.E. (2010). Executive dysfunction in traumatic brain injury: The effects of injury severity and effort on the Wisconsin Card Sorting Test. J. Clin. Exp. Neuropsychol. 32, 132–140. 10.1080/13803390902858874.

103. Miles, S., Howlett, C.A., Berryman, C., Nedeljkovic, M., Moseley, G.L., and Phillipou, A. (2021). Considerations for using the Wisconsin Card Sorting Test to assess cognitive flexibility. Behav. Res. Methods 53, 2083–2091.

105. Sarna, N., Dar, R., and Mazor, M. (2025). Biased and Inattentive Responding Drive Apparent Metacognitive Biases in Mental Health. Preprint at PsyArXiv. 10.31234/osf.io/fzwu8_v1.

105. Zorowitz, S., Solis, J., Niv, Y., and Bennett, D. (2023). Inattentive responding can induce spurious associations between task behaviour and symptom measures. Nat. Hum. Behav. 7, 1667–1681.

106. Baron-Cohen, S., Wheelwright, S., Skinner, R., Martin, J., and Clubley, E. (2001). The autism-spectrum quotient (AQ): evidence from Asperger syndrome/high-functioning autism, males and females, scientists and mathematicians. J. Autism Dev. Disord. 31, 5–17.

108. Vékony, T. (2021). Alternating Serial Reaction Time Task created with jsPsych (Version 1.0.0). Preprint.

109. Vékony, T. (2022). Card Sorting Task created with jsPsych (Version 1.0.1). Preprint.

110. de Leeuw, J.R. (2015). jsPsych: A JavaScript library for creating behavioral experiments in a Web browser. Behav. Res. Methods 47, 1–12. 10.3758/s13428-014-0458-y.

110. Fulop, F., Demeter, G., Honbolygó, F., and Csigo, K. (2024). Assessing obsessive-compulsive symptoms in a subclinical and clinical sample: the development of the Hungarian version of the OCI-R. Hungar Assoc Psychopharmacol 26, 144–152.

111. Anwyl-Irvine, A.L., Massonnié, J., Flitton, A., Kirkham, N., and Evershed, J.K. (2020). Gorilla in our midst: An online behavioral experiment builder. Behav. Res. Methods 52, 388–407. 10.3758/s13428-019-01237-x.

112. Vékony, T., Ambrus, G.G., Janacsek, K., and Nemeth, D. (2022). Cautious or causal? Key implicit sequence learning paradigms should not be overlooked when assessing the role of DLPFC (Commentary on Prutean et al.). Cortex 148, 222–226. 10.1016/j.cortex.2021.10.001.

114. R Core Team (2024). R: A language and environment for statistical computing (Version 4.4.2) [Computer software]. R Foundation for Statistical Computing. https://www.R-project.org/. Preprint.

114. Soetens, E., Melis, A., and Notebaert, W. (2004). Sequence learning and sequential effects. Psychological Research Psychologische Forschung 69, 124–137. 10.1007/s00426-003-0163-4.

116. Singmann, H., Bolker, B., Højsgaard, S., Fox, J., Lawrence, M.A., and Mertens, U. (2024). afex: Analysis of Factorial Experiments [Computer software]. Preprint.

116. Lenth, R. emmeans: Estimated Marginal Means, aka Least-Squares Means [Computer software]. https://CRAN.R-project.org/package=emmeans. Preprint.

