## Supplementary Information for "Obsessive-Compulsive Tendencies Shift the Balance Between Competitive Neurocognitive Functions"

**Supplementary Figures**

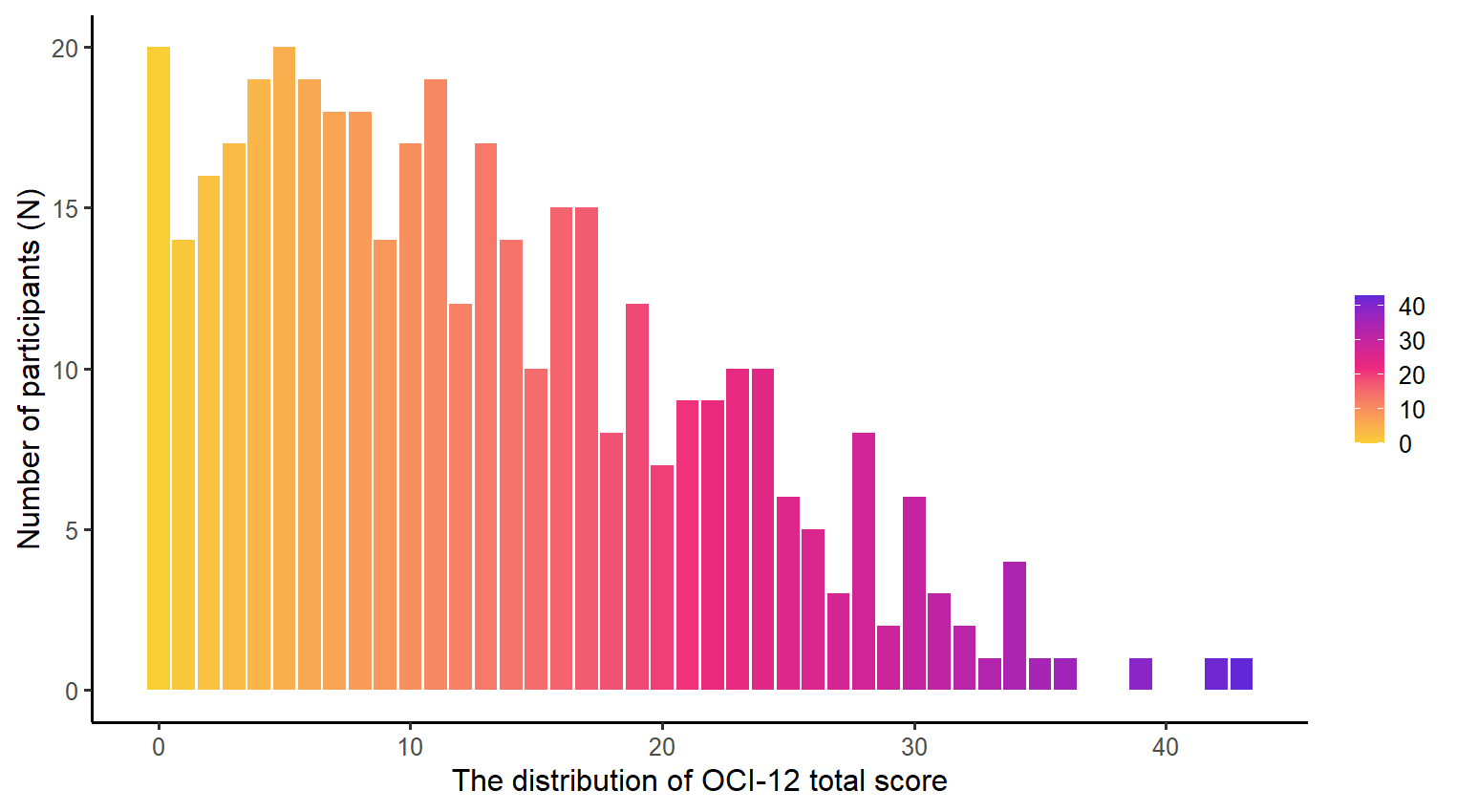

**Figure S1.** The distribution of the participants' OC tendencies as measured by the OCI-12 total score in the sample. The x-axis indicates the total OCI-12 score, while the y-axis indicates the number of participants. The color gradient transitions from yellow, through pink, to purple, reflecting increasing levels of OC tendencies.

**Supplementary Tables**

**Table S1. LMM predicting blockwise median RT, Related to Results**

|  | **Reaction Time** | | | | | |
| --- | --- | --- | --- | --- | --- | --- |
| *Terms* | *b* | *SE b* | *95% CI* | *t* | *df* | *p* |
| (Intercept) | 378.23 | 2.08 | 374.13 – 382.32 | 181.69 | 400.01 | **<0.001** |
| Time | -1.10 | 0.10 | -1.29 – -0.90 | -11.18 | 399.98 | **<0.001** |
| Triplet Type [High] | -2.94 | 0.23 | -3.38 – -2.49 | -13.01 | 11297.07 | **<0.001** |
| OCI-12 score | -0.42 | 0.23 | -0.88 – 0.04 | -1.77 | 400.01 | 0.077 |
| Perseverative error | 1.44 | 0.53 | 0.39 – 2.49 | 2.70 | 400.02 | **0.007** |
| Time × Triplet Type [High] | -0.17 | 0.05 | -0.27 – -0.07 | -3.23 | 11297.10 | **0.001** |
| Time × OCI-12 score | 0.00 | 0.01 | -0.02 – 0.03 | 0.39 | 399.88 | 0.699 |
| Triplet Type [High] × OCI-12 score | -0.00 | 0.03 | -0.05 – 0.05 | -0.09 | 11297.11 | 0.926 |
| Time × Perseverative error | -0.04 | 0.03 | -0.09 – 0.00 | -1.77 | 400.07 | 0.078 |
| Triplet Type [High] × Perseverative error | -0.04 | 0.06 | -0.15 – 0.08 | -0.66 | 11297.07 | 0.508 |
| OCI-12 score × Perseverative error | -0.03 | 0.06 | -0.14 – 0.08 | -0.52 | 400.01 | 0.600 |
| (Time × Triplet Type [High]) × OCI-12 score | 0.00 | 0.01 | -0.01 – 0.01 | 0.30 | 11297.18 | 0.763 |
| (Time × Triplet Type [High]) × Perseverative error | 0.01 | 0.01 | -0.01 – 0.04 | 0.96 | 11297.10 | 0.338 |
| (Time × OCI-12 score) × Perseverative error | 0.00 | 0.00 | -0.00 – 0.01 | 1.90 | 400.00 | 0.058 |
| (Triplet Type [High] × OCI-12 score) × Perseverative error | 0.01 | 0.01 | -0.00 – 0.02 | 1.13 | 11297.09 | 0.257 |
| (Time × Triplet Type [High] × OCI-12 score) × Perseverative error | -0.00 | 0.00 | -0.00 – 0.00 | -0.82 | 11297.15 | 0.412 |
| **Random Effects** | | | | | | |
| σ^2^ | 608.04 | | | | | |
| τ_00_ _Participant_ | 1703.21 | | | | | |
| τ_11_ _Participant.Time_ | 2.73 | | | | | |
| ρ_01_ _Participant_ | -0.30 | | | | | |
| ICC | 0.74 | | | | | |
| N _Participant_ | 404 | | | | | |
| Observations | 12113 | | | | | |
| Marginal R^2^ / Conditional R^2^ | 0.030 / 0.750 | | | | | |

Note. Statistical learning was evidenced by faster RTs for high-probability triplets compared to low-probability ones (main effect of Triplet Type). Reaction times improved throughout the task (main effect of Time). The interaction between Triplet Type and Block indicated that statistical learning (the difference between high- and low-probability triplets) became more pronounced as the task progressed.There was a statistically significant main effect of Perseverative Errors, indicating that more perseverative errors were associated with significantly slower reaction times. The absence of a significant interaction between Triplet Type and Perseverative Errors suggests that there is no antagonistic relationship between these factors.

**Table S2. LMM predicting blockwise mean accuracy, Related to Results**

|  | **Accuracy** | | | | | |
| --- | --- | --- | --- | --- | --- | --- |
| *Terms* | *b* | *SE b* | *95% CI* | *t* | *df* | *p* |
| (Intercept) | 90.90 | 0.17 | 90.56 – 91.24 | 525.51 | 400.00 | **<0.001** |
| Time | -0.15 | 0.02 | -0.18 – -0.11 | -7.67 | 400.22 | **<0.001** |
| Triplet Type [High] | 1.11 | 0.06 | 0.99 – 1.24 | 17.40 | 400.14 | **<0.001** |
| OCI-12 score | -0.00 | 0.02 | -0.04 – 0.04 | -0.03 | 399.97 | 0.976 |
| Perseverative error | -0.11 | 0.04 | -0.20 – -0.02 | -2.49 | 400.06 | **0.013** |
| Time × Triplet Type [High] | 0.08 | 0.01 | 0.05 – 0.11 | 5.76 | 10898.51 | **<0.001** |
| Time × OCI-12 score | -0.00 | 0.00 | -0.00 – 0.00 | -0.08 | 400.05 | 0.935 |
| Triplet Type [High] × OCI-12 score | 0.01 | 0.01 | -0.01 – 0.02 | 0.94 | 399.91 | 0.347 |
| Time × Perseverative error | -0.01 | 0.00 | -0.02 – -0.00 | -2.10 | 400.39 | **0.036** |
| Triplet Type [High] × Perseverative error | 0.03 | 0.02 | 0.00 – 0.07 | 2.12 | 400.52 | **0.035** |
| OCI-12 score × Perseverative error | -0.00 | 0.00 | -0.01 – 0.00 | -0.95 | 400.03 | 0.341 |
| (Time × Triplet Type [High])  × OCI-12 score | -0.00 | 0.00 | -0.00 – 0.00 | -0.99 | 10898.03 | 0.321 |
| (Time × Triplet Type [High])  × Perseverative error | -0.00 | 0.00 | -0.01 – 0.01 | -0.20 | 10899.01 | 0.845 |
| (Time × OCI-12 score) × Perseverative error | 0.00 | 0.00 | -0.00 – 0.00 | 1.52 | 400.26 | 0.130 |
| (Triplet Type [High] × OCI-12 score) ×  Perseverative error | -0.00 | 0.00 | -0.01 – -0.00 | -2.49 | 400.30 | **0.013** |
| (Time × Triplet Type [High]  × OCI-12 score) × Perseverative error | -0.00 | 0.00 | -0.00 – 0.00 | -1.35 | 10898.65 | 0.178 |
| **Random Effects** | | | | | | |
| σ^2^ | 41.20 | | | | | |
| τ_00_ _Participant_ | 10.53 | | | | | |
| τ_11_ _Participant.Time_ | 0.07 | | | | | |
| τ_11_ _Participant.Triplet Type [High]_ | 0.25 | | | | | |
| ρ_01_ | -0.05 | | | | | |
|  | -0.69 | | | | | |
| ICC | 0.23 | | | | | |
| N _Participant_ | 404 | | | | | |
| Observations | 12113 | | | | | |
| Marginal R^2^ / Conditional R^2^ | 0.036 / 0.255 | | | | | |

*Note.* Statistical learning was evidenced by higher accuracy for high-probability triplets compared to low-probability ones (main effect of Triplet Type). Accuracy declined over the course of the task (main effect of Block). The interaction between Triplet Type and Block indicated increasing statistical learning (the difference between high- and low-probability triplets) as the task progressed. A statistically significant main effect of Perseverative Errors suggested that higher perseverative error rates were associated with lower response accuracy, accompanied by a decline in visuomotor performance as the task progressed (Time × Perseverative error interaction). A significant interaction between Triplet Type and Perseverative Errors indicated an antagonistic relationship between statistical learning and cognitive flexibility. The significant Triplet Type × Perseverative Errors × OCI-R score interaction revealed that OC tendencies modulate this antagonistic relationship.

**Table S3. LMM predicting blockwise median RT, Related to Results**

| Effect | *df* | *F* | *p* |
| --- | --- | --- | --- |
| Time | 1, 399.99 | 120.03 | **< .001** |
| Triplet Type | 1, 11297.08 | 168.31 | **< .001** |
| OCI-12 score | 1, 400.00 | 2.44 | .119 |
| Trials to complete first category | 1, 400.00 | 1.26 | .263 |
| Time × Triplet Type | 1, 11297.13 | 11.08 | **< .001** |
| Time × OCI-12 score | 1, 399.88 | 0.28 | .595 |
| Triplet Type × OCI-12 score | 1, 11297.1 | 0.02 | .904 |
| Time × Trials to complete first category | 1, 399.83 | 0.23 | .635 |
| Triplet Type × Trials to complete first category | 1, 11297.04 | 0.20 | .652 |
| OCI-12 score × Trials to complete first category | 1, 400.00 | 0.02 | .887 |
| Time × Triplet Type × OCI-12 score | 1, 11297.15 | 0.07 | .790 |
| Time × Triplet Type × Trials to complete first category | 1, 11297.05 | 0.56 | .454 |
| Time × OCI-12 score × Trials to complete first category | 1, 399.80 | 0.06 | .801 |
| Triplet Type × OCI-12 score × Trials to complete first category | 1, 11297.04 | 0.23 | .632 |
| Time × Triplet Type × OCI-12 score × Trials to complete first category | 1, 11297.05 | 0.11 | .738 |

Note. The table summarizes all main effects and interactions. Degrees of freedom (*df)* correspond to Satterthwaite approximations. *F* = F-statistic; p = significance value. Statistically significant results (*p* < .05) are shown in bold. Significant main effects were followed up with post hoc comparisons: The main effect of Triplet Type (High vs. Low) was significant (*b* = -5.82, SE = 0.449, z = -12.973, *p* < .0001), indicating faster responses to high- than to low-probability triplets. Time was also significant (*b* = -1.07, SE = 0.0981, z = -10.956, *p* < .0001, 95% CI [-1.27, -0.882]), indicating that RT decreased as the task progressed. The Time × Triplet Type interaction was also significant (*b* = -0.346, SE = 0.104, z = -3.329, *p* = .0009), indicating a stronger decrease in RT for high-probability triplets over time. To depict the interaction, time was divided into three blocks (Block _5_, Block _10_, Block _15_). Pairwise comparisons showed significant High–Low differences at Block5 (*b* = -4.79, SE = 0.547, 95% CI [-5.86, -3.71]), Block_10_ (*b* = -6.52, SE = 0.495, 95% CI [-7.48, -5.55]), and Block_15_ (*b* = -8.24, SE = 0.855, 95% CI [-9.92, -6.57]). Sidak-adjusted comparisons indicated that the High–Low difference increased significantly across time: Block_5_–Block_10_ (b = 1.73, SE = 0.52, z = 3.329, p = .0026), Block_5_–Block_15_ (*b* = 3.46, SE = 1.04, z = 3.329, *p* = .0026), and Block_10_–Block_15_ (*b* = 1.73, SE = 0.52, z = 3.329, *p* = .0026). These results indicate a gradual increase in statistical learning across the task.

**Table S4. LMM predicting blockwise mean accuracy, Related to Results**

| Effect | *df* | *F* | *p* |
| --- | --- | --- | --- |
| Time | 1, 400.25 | 55.48 | **< .001** |
| Triplet Type | 1, 399.97 | 288.22 | **< .001** |
| OCI-12 score | 1, 399.96 | 0.57 | .450 |
| Trials to complete first category | 1, 399.94 | 0.22 | .639 |
| Time × Triplet Type | 1, 10898.42 | 31.05 | **< .001** |
| Time × OCI-12 score | 1, 400.08 | 0.05 | .824 |
| Triplet Type × OCI-12 score | 1, 399.77 | 1.10 | .295 |
| Time × Trials to complete first category | 1, 399.98 | 1.55 | .214 |
| Triplet Type × Trials to complete first category | 1, 399.66 | 0.17 | .680 |
| OCI-12 score × Trials to complete first category | 1, 399.94 | 2.18 | .141 |
| Time × Triplet Type × OCI-12 score | 1, 10897.91 | 1.3 | .255 |
| Time × Triplet Type × Trials to complete first category | 1, 10897.59 | 0.58 | .446 |
| Time × OCI-12 score × Trials to complete first category | 1, 399.94 | 0.52 | .470 |
| Triplet Type × OCI-12 score × Trials to complete first category | 1, 399.62 | 4.58 | **.033** |
| Time × Triplet Type × OCI-12 score × Trials to complete first category | 1, 10897.47 | 0.77 | .379 |

Note. The table summarizes all main effects and interactions. Degrees of freedom *(df)* correspond to Satterthwaite approximations. *F* = F-statistic; *p* = significance value. Statistically significant results (p < .05) are shown in bold. Significant main effects were followed up with post hoc comparisons:. The main effect of Triplet Type (High vs. Low) was significant (*b* = 2.17, SE = 0.128, z = 16.977, *p* < .0001), indicating higher accuracy for high- than for low-probability triplets. The linear trend of Time was also significant (*b* = -0.141, SE = 0.0189, z = -7.448, *p* < .0001, 95% CI [-0.178, -0.104]), indicating that accuracy decreased as the task progressed. To depict the Time × Triplet Type interaction, time was segmented into three blocks (Block_5_, Block_10_, Block_15_). Pairwise comparisons showed significant High–Low differences at Block_5_ (*b* = 1.72, SE = 0.151, 95% CI [1.42, 2.01]), Block_10_ (*b* = 2.47, SE = 0.139, 95% CI [2.20, 2.74]), and Block1_5_ (*b* = 3.22, SE = 0.228, 95% CI [2.78, 3.67]). Sidak-adjusted comparisons indicated that the High–Low difference increased significantly across time: Block_5_–Block_10_ (b = -0.754, SE = 0.135, z = -5.572, *p* < .0001), Block_5_–Block_15_ (*b* = -1.507, SE = 0.270, z = -5.572, *p* < .0001), and Block10–Block15 (*b* = -0.754, SE = 0.135, z = -5.572, *p* < .0001), indicating increasing statistical learning over time. The significant three-way interaction, we examined the simple effect of triplet type (high- vs. low-probability triplets) at −1 SD, mean, and +1 SD levels of Trials to complete the first category and OCI-12 scores. Please note that, we used the OCI-12 score and the Trials to complete the first category as a continuous variable for all our analyses; we only depicted results for post-hoc analysis. Among individuals with low OC tendencies, the SL performance decreased as the number of trials to complete the first category increased. When participants completed the first category quickly, the statistical learning was largest, *b* = 2.42, SE = 0.26, z = 9.17, 95% CI [1.90, 2.94], *p* < .001. At the mean level of trials, the effect was smaller, *b* = 2.03, SE = 0.18, z = 11.16, 95% CI [1.68, 2.39], p < .001, and it was smallest at later trials, *b* = 1.65, SE = 0.30, z = 5.48, 95% CI [1.06, 2.24], *p* < .001. At a medium OC tendencies, statistical remained relatively stable across levels of trials (b_earlier trials_ = 2.22; *b_mean_ trials* = 2.17;: *_blater trials_* = 2.11; all *ps* < .001), suggesting little modulation of statistical learning by cognitive flexibility. In contrast, among individuals with higher OC tendencies, the pattern reversed. The triplet effect increased as trials to complete the first category increased. At earlier trials, the effect was *b* = 2.02, SE = 0.26, z = 7.86, 95% CI [1.52, 2.53], *p* < .001; at mean trials, *b* = 2.30, SE = 0.18, z = 12.75, 95% CI [1.95, 2.66], *p* < .001; and at later trials, the effect was strongest, *b* = 2.58, SE = 0.26, z = 9.84, 95% CI [2.07, 3.10*], p* < .001. This indicates that among individuals with higher OC tendencies, poorer CST performance was associated with enhanced statistical learning.

**Table S5. LMM predicting blockwise median RT, Related to Results**

| Effect | *df* | *F* | *p* |
| --- | --- | --- | --- |
| Time | 1, 399.95 | 127.73 | **< .001** |
| Triplet Type | 1, 11297.06 | 170.45 | **< .001** |
| OCI-12 score | 1, 400.00 | 4.66 | **.032** |
| Total correct | 1, 400.01 | 12.55 | **< .001** |
| Time × Triplet Type | 1, 11297.08 | 9.73 | **.002** |
| Time × OCI-12 score | 1, 399.87 | 0.25 | .615 |
| Triplet Type × OCI-12 score | 1, 11297.0917 | 0.00 | .960 |
| Time × Total correct | 1, 400.01 | 2.21 | .138 |
| Triplet Type × Total correct | 1, 11297.10 | 0.27 | .607 |
| OCI-12 score × Total correct | 1, 400.00 | 0.10 | .750 |
| Time × Triplet Type × OCI-12 score | 1, 11297.15 | 0.15 | .698 |
| Time × Triplet Type × Total correct | 1, 11297.17 | 0.00 | .952 |
| Time × OCI-12 score × Total correct | 1, 399.96 | 6.02 | **.015** |
| Triplet Type × OCI-12 score × Total correct | 1, 11297.14 | 2.55 | .110 |
| Time × Triplet Type × OCI-12 score × Total correct | 1, 11297.23 | 1.36 | .244 |

Note.The table summarizes all main effects and interactions. Degrees of freedom (*df)* correspond to Satterthwaite approximations. *F* = F-statistic; p = significance value. Statistically significant results (*p* < .05) are shown in bold. Significant main effects were followed up with post hoc comparisons: The main effect of Triplet Type (High vs. Low) was significant (*b* = -5.93, SE = 0.454, z = -13.056, *p* < .0001), indicating faster responses to high- than to low-probability triplets. The linear trend of Time was also significant (*b* = -1.11, SE = 0.0984, z = -11.302, *p* < .0001, 95% CI [-1.31, -0.92]), showing that RT decreased as the task progressed. The Time × Triplet Type interaction was significant (b = -0.328, SE = 0.105, z = -3.120, p = .0018), indicating a stronger decrease in RT for high-probability triplets over time. To illustrate this interaction, time was depicted at three blocks (Block_5_, Block_10_, Block_15_).Pairwise comparisons showed significant High–Low differences at Block_5_ (*b* = -4.95, SE = 0.553, 95% CI [-6.03, -3.86]), Block_10_ (*b* = -6.59, SE = 0.501, 95% CI [-7.57, -5.60]), and Block_15_ (*b* = -8.23, SE = 0.865, 95% CI [-9.92, -6.53]). Sidak-adjusted comparisons indicated that the High–Low difference increased significantly across time: Block_5_–Block_10_ (*b* = 1.64, SE = 0.526, z = 3.120, *p* = .0054), Block_5_–Block_15_ (*b* = 3.28, SE = 1.050, z = 3.120, *p* = .0054), and Block_10_–Block_15_ (b = 1.64, SE = 0.526, z = 3.120, p = .0054), suggesting a gradual increase in statistical learning during the task. Furthermore, there was a significant negative main effect of OCI-12 on median RT, *b* = −0.498, SE = 0.231, 95% CI [−0.95, −0.046], indicating that higher OCI-12 scores were associated with faster reaction times. Over time, the number of correct responses, and the leve lof OCI tendencies, RT decreased with later blocks (learning effect; bs ~380 ms in early blocks vs. ~376 ms in late blocks), were slightly faster for correct responses (~2 ms difference), and were faster for participants with higher OCI-12 scores (~2–3 ms faster from low to high OCI-12), with minimal evidence for a strong three-way interaction (*p* < .001).

**Table S6. LMM predicting blockwise mean accuracy, Related to Results**

| Effect | *df* | *F* | *p* |
| --- | --- | --- | --- |
| Time | 1, 400.20 | 60.15 | **< .001** |
| Triplet Type | 1, 400.09 | 294.22 | **< .001** |
| OCI-12 score | 1, 399.97 | 0.01 | .906 |
| Total correct | 1, 400.02 | 4.01 | **.046** |
| Time × Triplet Type | 1, 10,898.40 | 34.78 | **< .001** |
| Time × OCI-12 score | 1, 400.05 | 0.00 | .988 |
| Triplet Type × OCI-12 score | 1, 399.90 | 0.30 | .582 |
| Time × Total correct | 1, 400.30 | 2.62 | .106 |
| Triplet Type × Total correct | 1, 400.20 | 4.58 | **.033** |
| OCI-12 score × Total correct | 1, 400.00 | 0.79 | .375 |
| Time × Triplet Type × OCI-12 score | 1, 10897.99 | 1.19 | .275 |
| Time × Triplet Type × Total correct | 1, 10898.73 | 0.04 | .847 |
| Time × OCI-12 score × Total correct | 1, 400.22 | 3.88 | **.050** |
| Triplet Type × OCI-12 score × Total correct | 1, 400.11 | 2.07 | .151 |
| Time × Triplet Type × OCI-12 score × Total correct | 1, 10898.51 | 4.31 | **.038** |

Note. The table summarizes all main effects and interactions. Degrees of freedom (*df)* correspond to Satterthwaite approximations. *F* = F-statistic; p = significance value. Statistically significant results (*p* < .05) are shown in bold. Significant main effects were followed up with post hoc comparisons: The main effect of Triplet Type (High vs. Low) was significant (*b* = 2.21, SE = 0.129, z = 17.153, *p* < .0001), indicating higher accuracy for high- than for low-probability triplets. The linear trend of Time was also significant (*b* = −0.148, SE = 0.019, z = −7.755, *p* < .0001, 95% CI [−0.185, −0.110]), showing that accuracy decreased as the task progressed. The Time × Triplet Type interaction indicated that the High–Low difference increased across time. To illustrate this interaction, time was depicted at three blocks (Block_5_, Block_10_, Block_15_). Pairwise comparisons at three representative blocks showed significant High–Low differences at Block _5_ (*b* = 1.73, SE = 0.153, 95% CI [1.43, 2.03]), Block _10_ (*b* = 2.54, SE = 0.140, 95% CI [2.26, 2.81]), and Block _15_ (*b* = 3.34, SE = 0.231, 95% CI [2.89, 3.80]). Šidák-adjusted comparisons confirmed that the High–Low difference increased significantly across blocks: Block_5_–Block_10_ (*b* = 0.807, SE = 0.137, z = −5.898, *p* < .0001), Block_5_–Block_15_ (b = 1.614, SE = 0.274, z = −5.898, *p* < .0001), and Block_10_–Block_15_ (*b* = 0.807, SE = 0.137, z = −5.898, *p* < .0001). The overall linear trend for correct performance was significant (*b* = 0.0371, SE = 0.0185, z = 2.002, 95% CI [0.0008, 0.0735], *p* = 0.045), indicating that higher numbers of correct responses were associated with more accurate responses. The Triplet Type x Correct responses interaction revealed decreasing statistical learning with increased rate of correct responses (b _Less correct_ = 2.24%, 95% CI [1.99, 2.50]), mean (b _Mean correct_ = 2.21%, 95% CI [1.96, 2.47]), and (b _More correct_ = 2.19%, 95% CI [1.93, 2.44]) (all p < .0001). The four-way interaction indicated that the block-related accuracy decline was driven exclusively by low-probability (L) triplets, whereas high-probability (H) triplets remained stable across blocks. This pattern was consistent across OCI-12 and correctness levels, suggesting limited moderation of temporal learning dynamics by OC tendencies or response accuracy.

**Table S7. LMM predicting blockwise median RT, Related to Results**

| Effect | ***df*** | ***F*** | ***p*** |
| --- | --- | --- | --- |
| Time | 399.9713 | 124.248 | < .001 |
| Triplet Type | 11,297.0703 | 169.367 | < .001 |
| OCI-R score | 400.0016 | 6.314 | .012 |
| Perseverative error | 400.0107 | 7.197 | .008 |
| Time × Triplet Type | 11,297.1001 | 10.343 | .001 |
| Time × OCI-R score | 399.9023 | 1.047 | .307 |
| Triplet Type × OCI-R score | 11,297.1190 | 0.003 | .959 |
| Time × Perseverative error | 400.0583 | 3.242 | .073 |
| Triplet Type × Perseverative error | 11,297.0705 | 0.497 | .481 |
| OCI-R score × Perseverative error | 400.0054 | 0.036 | .851 |
| Time × Triplet Type × OCI-R score | 11,297.1940 | 0.000 | .985 |
| Time × Triplet Type × Perseverative error | 11,297.1004 | 0.990 | .320 |
| Time × OCI-R score × Perseverative error | 399.9603 | 2.503 | .114 |
| Triplet Type × OCI-R score × Perseverative error | 11,297.1020 | 1.491 | .222 |
| Time × Triplet Type × OCI-R score × Perseverative error | 11,297.1611 | 0.653 | .419 |

Note. The table summarizes all main effects and interactions. Degrees of freedom (*df)* correspond to Satterthwaite approximations. *F* = F-statistic; p = significance value. Statistically significant results (*p* < .05) are shown in bold. Significant main effects were followed up with post hoc comparisons: The main effect of Triplet Type (High vs. Low) was significant (b = −5.89, SE = 0.452, z = −13.014, p < .0001), indicating faster responses to high-probability triplets. The main effect of Time indicated a decrease in RTs throughout the task, demonstrating changes in visuomotor performance over time (b = -1.09, 95% CI =[-1.29, -.90). The Time x Triplet Type interaction was examined at three representative blocks (Block _5,_ Block _10_, Block _15_). Pairwise comparisons showed significant statistical learning Block_5_ (b = −4.88, SE = 0.551, 95% CI [−5.95, −3.80]), Block_10_ (b = −6.56, SE = 0.498, 95% CI [−7.54, −5.58]), and Block_15_ (b = −8.24, SE = 0.861, 95% CI [−9.93, −6.55]), indicating that the High–Low RT advantage increased across time. Šidák-adjusted interaction contrasts confirmed that the statistical increased significantly over blocks: Block5–Block10 (b = 1.68, SE = 0.523, z = 3.216, p = .0039), Block_5_–Block_15_ (b = 3.37, SE = 1.050, z = 3.216, p = .0039), and Block_10_–Block_15_ (b = 1.68, SE = 0.523, z = .0039), suggesting a gradual strengthening of learning-related performance improvement over time.main effect of Perseverative error indicating that weaker cognitive flexibility (more perseverative errors) was associated with significantly slower reaction times (b = 1.31, 95% CI =[0.38, 2.24]). ). Furthermore, the main effect of OCI-R score was statistically significant indicating faster reaction times with higher levels of OC tendencies (b = -0.44, 95% CI =[-0.74, -0.12]).

**Table S8. LMM predicting blockwise median RT, Related to Results**

|  | **Reaction Time** | | | | | |
| --- | --- | --- | --- | --- | --- | --- |
| *Terms* | *b* | *SE b* | *95% CI* | *t* | *df* | *p* |
| (Intercept) | 378.04 | 2.08 | 373.95 – 382.12 | 181.93 | 400.01 | **<0.001** |
| Time | -1.09 | 0.10 | -1.29 – -0.90 | -11.15 | 399.97 | **<0.001** |
| Triplet Type [High] | -2.94 | 0.23 | -3.39 – -2.50 | -13.01 | 11297.07 | **<0.001** |
| OCI-R score | -0.45 | 0.18 | -0.80 – -0.10 | -2.51 | 400.00 | **0.012** |
| Perseverative error | 1.43 | 0.53 | 0.38 – 2.48 | 2.68 | 400.01 | **0.008** |
| Time × Triplet Type [High] | -0.17 | 0.05 | -0.27 – -0.07 | -3.22 | 11297.10 | **0.001** |
| Time × OCI-R score | 0.01 | 0.01 | -0.01 – 0.03 | 1.02 | 399.90 | 0.307 |
| Triplet Type [High] × OCI-R score | 0.00 | 0.02 | -0.04 – 0.04 | 0.05 | 11297.12 | 0.959 |
| Time × Perseverative error | -0.05 | 0.03 | -0.09 – 0.00 | -1.80 | 400.06 | 0.073 |
| Triplet Type [High] × Perseverative error | -0.04 | 0.06 | -0.15 – 0.07 | -0.71 | 11297.07 | 0.481 |
| OCI-R score × Perseverative error | 0.01 | 0.04 | -0.07 – 0.09 | 0.19 | 400.01 | 0.851 |
| (Time × Triplet Type [High]) × OCI-R score | 0.00 | 0.00 | -0.01 – 0.01 | 0.02 | 11297.19 | 0.985 |
| (Time × Triplet Type [High]) × Perseverative error | 0.01 | 0.01 | -0.01 – 0.04 | 1.00 | 11297.10 | 0.320 |
| (Time × OCI-R score) × Perseverative error | 0.00 | 0.00 | -0.00 – 0.01 | 1.58 | 399.96 | 0.114 |
| (Triplet Type [High] × OCI-R score) × Perseverative error | 0.01 | 0.00 | -0.00 – 0.01 | 1.22 | 11297.10 | 0.222 |
| (Time × Triplet Type [High] × OCI-R score) × Perseverative error | -0.00 | 0.00 | -0.00 – 0.00 | -0.81 | 11297.16 | 0.419 |
| **Random Effects** | | | | | | |
| σ^2^ | 608.03 | | | | | |
| τ_00_ _Participant_ | 1692.42 | | | | | |
| τ_11_ _Participant.Time_ | 2.73 | | | | | |
| ρ_01_ _Participant_ | -0.30 | | | | | |
| ICC | 0.74 | | | | | |
| N _Participant_ | 404 | | | | | |
| Observations | 12113 | | | | | |
| Marginal R^2^ / Conditional R^2^ | 0.034 / 0.750 | | | | | |

Note. Statistical learning was evidenced by faster RTs for high-probability triplets compared to low-probability ones (main effect of Triplet Type). Reaction times improved throughout the task (main effect of Time). The interaction between Triplet Type and Block indicated that statistical learning (the difference between high- and low-probability triplets) became more pronounced as the task progressed.There was a statistically significant main effect of Perseverative Errors, indicating that more perseverative errors were associated with significantly slower reaction times. The absence of a significant interaction between Triplet Type and Perseverative Errors suggests that there is no antagonistic relationship between these factors.

**Table S9. LMM predicting blockwise mean accuracy, Related to Results**

| *Effect* | *df* | *F* | *p* |
| --- | --- | --- | --- |
| Time | 1, 400.22 | 56.95 | **< .001** |
| Triplet Type | 1, 400.12 | 300.37 | **< .001** |
| OCI-R score | 1, 399.97 | 0.17 | .685 |
| Perseverative error | 1, 400.06 | 5.89 | .**016** |
| Time × Triplet Type | 1, 10898.47 | 33.1 | **< .001** |
| Time × OCI-R score | 1, 400.10 | 0.25 | .616 |
| Triplet Type × OCI-R score | 1, 399.92 | 0.32 | .571 |
| Time × Perseverative error | 1, 400.38 | 3.99 | **.046** |
| Triplet Type × Perseverative error | 1, 400.47 | 4.42 | **.036** |
| OCI-R score × Perseverative error | 1, 400.01 | 0.88 | .349 |
| Time × Triplet Type × OCI-R score | 1, 10898.15 | 0.72 | .395 |
| Time × Triplet Type × Perseverative error | 1, 10898.93 | 0.05 | .829 |
| Time × OCI-R score × Perseverative error | 1, 400.21 | 0.45 | .502 |
| Triplet Type × OCI-R score × Perseverative error | 1, 400.15 | 4.92 | **.027** |
| Time × Triplet Type × OCI-R score × Perseverative error | 1, 10898.44 | 1.6 | .207 |

Note. The table summarizes all main effects and interactions. Degrees of freedom (*df)* correspond to Satterthwaite approximations. *F* = F-statistic; p = significance value. Statistically significant results (*p* < .05) are shown in bold. Significant main effects were followed up with post hoc comparisons: The main effect of Triplet Type was significant (b = 2.22, SE = 0.128, z = 17.331, p < .0001), indicating higher accuracy for high- than for low-probability triplets. Pairwise comparisons showed a significant High–Low difference (b = −5.89, SE = 0.452, z = −13.014, p < .0001). The High–Low accuracy difference increased across time blocks, with significant differences at Block5 (b = 1.75, SE = 0.152, 95% CI [1.45, 2.05]), Block10 (b = 2.54, SE = 0.139, 95% CI [2.26, 2.81]), and Block15 (b = 3.32, SE = 0.230, 95% CI [2.87, 3.77]). Šidák-adjusted interaction contrasts indicated that the High–Low difference increased significantly across blocks (Block5–Block10: b = 0.784, SE = 0.136, z = −5.753, p < .0001; Block5–Block15: b = 1.568, SE = 0.273, z = −5.753, p < .0001; Block10–Block15: b = 0.784, SE = 0.136, z = −5.753, p < .0001). Perseverative error tendency showed a significant negative association with accuracy (trend estimate = −0.108, SE = 0.044, asymp. 95% CI [−0.195, −0.021]).Triplet Type * Perseverative error interaction demonstrated an antagonistic relationship between statistical learning and cognitive flexibility, with greater statistical learning linked to reduced cognitive flexibility (higher number of perseverative errors). (*b_Low Perseverative errors_* = 1.95, 95% CI =[1.59, 2.30]; *b_Mean Perseverative errors_*  = 2.22, 95% CI =[1.97, 2.47]; *b_High Perseverative errors_*  = 2.50, 95% CI =[2.13, 2.86]).Triplet Type * Perseverative error * OCI-R score (F(1, 400.15) = 4.92, p = .027) highlighted the modulating role of OC tendencies in the antagonistic relationship between statistical learning and cognitive flexibility (Figure. 4). The typical antagonistic relationship between cognitive flexibility and statistical learning, where poorer flexibility is associated with enhanced statistical learning, was observed in individuals with lower and medium OC tendencies. Please note that, we used the OCI-R score as a continuous variable for all our analyses, for post-hoc analysis, we depicted results at lower OC tendencies (Mean OCI-R score - SD), medium OC tendencies (Mean OCI-R score) and higher OC tendencies (Mean OCI-R score + SD), based on standard deviations from the mean. For lower OC tendencies, statistical learning (High – Low difference) was smallest at a low number of perseverative errors (_blow perseverative errors_ = 1.60, 95% CI [1.10, 2.11]), increased at mean perseverative errors (b _Mean Perseverative errors_ = 2.15, 95% CI [1.79, 2.50]), and was largest at high perseverative errors (b _High Perseverative errors_ = 2.69, 95% CI [2.16, 3.22]). For medium OC tendencies, the corresponding differences were _bLow Perseverative errors_ = 1.95, 95% CI [1.59, 2.30]; b _Mean Perseverative errors_ = 2.22, 95% CI [1.97, 2.47]; and _bHigh Perseverative errors_ = 2.50, 95% CI [2.13, 2.86]. However, this relationship weakened in individuals with high OC tendencies: b _Low Perseverative errors_ = 2.29, 95% CI [1.76, 2.82]; b_Mean Perseverative_ _errors_ = 2.30, 95% CI [1.93, 2.66]; and b_High Perseverative errors_ = 2.30, 95% CI [1.88, 2.73]. Overall, the difference between high and low triplet conditions increased with higher perseverative errors under lower OC tendencies but remained relatively stable when OC tendencies were high. With higher OC tendencies, statistical learning was not enhanced in individuals with poor cognitive flexibility; instead, learning remained at a modest level, regardless of their cognitive flexibility performance.

**Table S10. LMM predicting blockwise mean accuracy, Related to Results**

|  | **Accuracy** | | | | | |
| --- | --- | --- | --- | --- | --- | --- |
| *Terms* | *b* | *SE b* | *95% CI* | *t* | *df* | *p* |
| (Intercept) | 90.90 | 0.17 | 90.56 – 91.24 | 524.99 | 400.01 | **<0.001** |
| Time | -0.14 | 0.02 | -0.18 – -0.12 | -7.57 | 400.22 | **<0.001** |
| Triplet Type [High] | 1.11 | 0.06 | 0.99 – 1.2 | 17.33 | 400.12 | **<0.001** |
| OCI-R score | -0.01 | 0.01 | -0.04 – 0.02 | -0.41 | 399.97 | 0.685 |
| Perseverative error | -0.11 | 0.04 | -0.2 – -0.02 | -2.43 | 400.06 | **0.016** |
| Time × Triplet Type [High] | 0.08 | 0.01 | 0.05 – 0.11 | 5.75 | 10898.47 | **<0.001** |
| Time × OCI-R score | 0.00 | 0.00 | -0.00 – 0.00 | 0.50 | 400.10 | 0.616 |
| Triplet Type [High] × OCI-R score | 0.00 | 0.01 | -0.01 – 0.01 | 0.57 | 399.92 | 0.571 |
| Time × Perseverative error | -0.01 | 0.00 | -0.02 – -0.00 | -1.99 | 400.38 | **0.046** |
| Triplet Type [High] × Perseverative error | 0.03 | 0.02 | 0.00 – 0.07 | 2.10 | 400.47 | **0.036** |
| OCI-R score × Perseverative error | -0.00 | 0.00 | -0.01 – 0.00 | -0.94 | 400.01 | 0.349 |
| (Time × Triplet Type [High]) × OCI-R score | -0.00 | 0.00 | -0.00 – 0.00 | -0.85 | 10898.15 | 0.395 |
| (Time × Triplet Type [High]) × Perseverative error | -0.00 | 0.00 | -0.01 – 0.01 | -0.22 | 10898.93 | 0.829 |
| (Time × OCI-R score) × Perseverative error | 0.00 | 0.00 | -0.00 – 0.00 | 0.67 | 400.21 | 0.502 |
| (Triplet Type [High] × OCI-R score) × Perseverative error | -0.00 | 0.00 | -0.01 – -0.00 | -2.22 | 400.15 | **0.027** |
| (Time × Triplet Type [High] × OCI-R score) × Perseverative error | -0.00 | 0.00 | -0.00 – 0.00 | -1.26 | 10898.44 | 0.207 |
| **Random Effects** | | | | | | |
| σ^2^ | 41.21 | | | | | |
| τ_00_ _Participant_ | 10.52 | | | | | |
| τ_11_ _Participant.Time_ | 0.07 | | | | | |
| τ_11_ _Participant.Time.Triplet Type_ | 0.26 | | | | | |
| ρ_01_ | -0.05 | | | | | |
|  | -0.68 | | | | | |
| ICC | 0.23 | | | | | |
| N _Participant_ | 404 | | | | | |
| Observations | 12113 | | | | | |
| Marginal R^2^ / Conditional R^2^ | 0.036 / 0.255 | | | | | |

*Note.* Statistical learning was evidenced by higher accuracy for high-probability triplets compared to low-probability ones (main effect of Triplet Type). Accuracy declined over the course of the task (main effect of Time). The interaction between Triplet Type and Time indicated increasing statistical learning (the difference between high- and low-probability triplets) as the task progressed.A statistically significant main effect of Perseverative Errors suggested that higher perseverative error rates were associated with lower response accuracy, accompanied by a decline in visuomotor performance as the task progressed. A significant Triplet Type and Perseverative Errors interaction indicated an antagonistic relationship between statistical learning and cognitive flexibility. The significant Triplet Type × Perseverative Errors × OCI-R score interaction revealed that OC tendencies modulate this antagonistic relationship.
